# Novel chromosome-length genome assemblies of three distinct subspecies of pine marten, sable, and yellow-throated marten (genus *Martes*, family Mustelidae)

**DOI:** 10.1101/2025.09.22.677678

**Authors:** Andrey A. Tomarovsky, Ruqayya Khan, Olga Dudchenko, Violetta R. Beklemisheva, Polina L. Perelman, Azamat A. Totikov, Natalia A. Serdyukova, Tatiana M. Bulyonkova, Maria Pobedintseva, Alexei V. Abramov, David Weisz, Aliya Yakupova, Anna Zhuk, Alexander S. Graphodatsky, Roger Powell, Erez Lieberman Aiden, Klaus-Peter Koepfli, Sergei Kliver

**Affiliations:** Laboratory of Diversity and Evolution of Genomes, Institute of Molecular and Cellular Biology SB RAS, 8/2 Acad. Lavrentiev ave., Novosibirsk, 630090, Russia; Department of Natural Sciences, Novosibirsk State University, 1 Pirogova str., Novosibirsk, 630090, Russia; The Center for Genome Architecture, Department of Molecular and Human Genetics, Baylor College of Medicine, Houston, TX 77030, USA; The Center for Theoretical Biological Physics, Rice University, TX 77005, USA; Laboratory of System Dynamics, A. P. Ershov Institute of Informatics Systems SB RAS, 6 Acad. Lavrentjev ave., Novosibirsk, 630090, Russia; Laboratory for Theriology, Zoological Institute RAS, 1 Universitetskaya emb., St. Petersburg, 199034, Russia; Division of Evolutionary Biology, Ludwig-Maximilians-Universität, 2, Großhaderner str, Planegg, 82152, Germany; Microevolution and Biodiversity, Max Planck Institute for Biological Intelligence, Eberhard-Gwinner-Straße, Seewiesen, 82319, Germany; Institute of Applied Computer Science, ITMO University, 197101 St. Petersburg, Russia; Laboratory of Amyloid Biology, St. Petersburg State University, 199034 St. Petersburg, Russia; North Carolina State University; Smithsonian-Mason School of Conservation, 1500 Remount Road, Front Royal, VA 22630, USA; Center for Evolutionary Hologenomics, The Globe Institute, The University of Copenhagen, 5A, Oester Farimagsgade, Copenhagen, 1353, Denmark

## Abstract

The genus *Martes* consists of medium-sized carnivores within the family Mustelidae that are commonly known as martens, many of which exhibit extensive geographic variation and taxonomic uncertainty. Here, we report chromosome-length genome assemblies for three subspecies, each representing a different marten species: the Tobol sable (*M. zibellina zibellina*), the Ural pine marten (*M. martes uralensis*), and the Far East yellow-throated marten (*M. flavigula aterrima*). Using linked-read sequencing and Hi-C scaffolding, we generated assemblies with total lengths of 2.39-2.45 Gbp, N50 values of 137-145 Mbp, and high BUSCO scores (93.6-96.4%). We identified 19 chromosomal scaffolds for sable and pine marten, and 20 for yellow-throated marten, which agrees with the known karyotypes of these species (2n=38 and 2n=40, respectively). Annotation predicted ~20,000 protein-coding genes per genome, of which >90% were assigned functional names. Repeats encompass 36.9-40.4% of the assemblies, with a prevalence of LINEs and SINEs, and is conservative across the genus. Synteny analysis of our generated and available marten genome assemblies revealed assembly artifacts in previously published assemblies, which we confirmed through investigation of Hi-C contact maps. Among other rearrangements, we verified a sable-specific inversion on chromosome 11 using the published cytogenetic data. Our assemblies broaden the genomic resources available for *Martes*, extending coverage to geographically distant and taxonomically significant subspecies. Together, they provide a robust framework for assessing intraspecific genetic diversity, identifying signatures of hybridization, and refining the complex taxonomy of the genus. Beyond conservation and evolutionary applications, these references will facilitate comparative genomics across Mustelidae and other carnivorans.

## Introduction

The genus *Martes* (family Mustelidae) comprises medium-sized carnivores, distributed mainly across the Holarctic region [1]. According to current taxonomy, *Martes* is divided into two subgenera: *Martes* (six species) and *Charronia* (*M. flavigula* and *M. gwatkinsii*) [2, 3]. Most of the species are distributed across the Palearctic, for example, the sable (*M. zibellina*), the pine marten (*M. martes*), the stone marten (*M. foina*), and the Japanese marten (*M. melampus*) [4–7]. In the Nearctic, the American marten (*M. americana*) and Pacific marten (*M. caurina*) are found [8, 9], while the Nilgiri yellow-throated marten (*M. gwatkinsii*) and yellow-throated marten (*M. flavigula*) inhabit the Indomalayan region, the latter also extending into the northeastern Palearctic (Figure **1**) [10, 11]. Most of these species occupy broad ranges and show substantial intraspecific diversity, which is reflected by the large number of described subspecies (Supplementary Table ST1). *M. zibellina* is known to have the highest subspecific diversity, as up to 17 subspecies are recognized [1, 12]. A considerable number of subspecies have also been described for *M. flavigula* (about 10) [13, 14], *M. martes* (10) [15], and *M. foina* (11) [1]. Other species harbor fewer subspecies: up to six each in *M. americana* and *M. caurina* [16], three in *M. melampus* [17, 18], whereas the narrowly distributed *M. gwatkinsii* is currently regarded as monotypic. However, these subspecies classifications should be considered preliminary, as they are primarily based on phenotypic and morphometric traits, and only a few of them have been supported by genetic data [19–22]. Many described forms have uncertain taxonomic status and may eventually be synonymized, merged with other subspecies, or, conversely, recognized as distinct species, which was the case for *M. americana* and *M. caurina*, the latter being previously regarded as belonging to *M. americana* [23, 9]. This problem is especially evident in *M. flavigula*. Numerous morphological and geographic forms have been described, and repeated attempts to organize them have often resulted in proposals to elevate certain populations to species rank. For example, some authors have suggested recognizing populations from the Russian Far East and Indochina as distinct species, either within the subgenus *Charronia* or even by assigning *Charronia* to full genus status within Guloninae [24, 25]. Such debates highlight the complexity of intraspecific structure in *M. flavigula* and the challenges of its taxonomic interpretation, making it one of the most problematic species within the genus.

**Figure 1.**
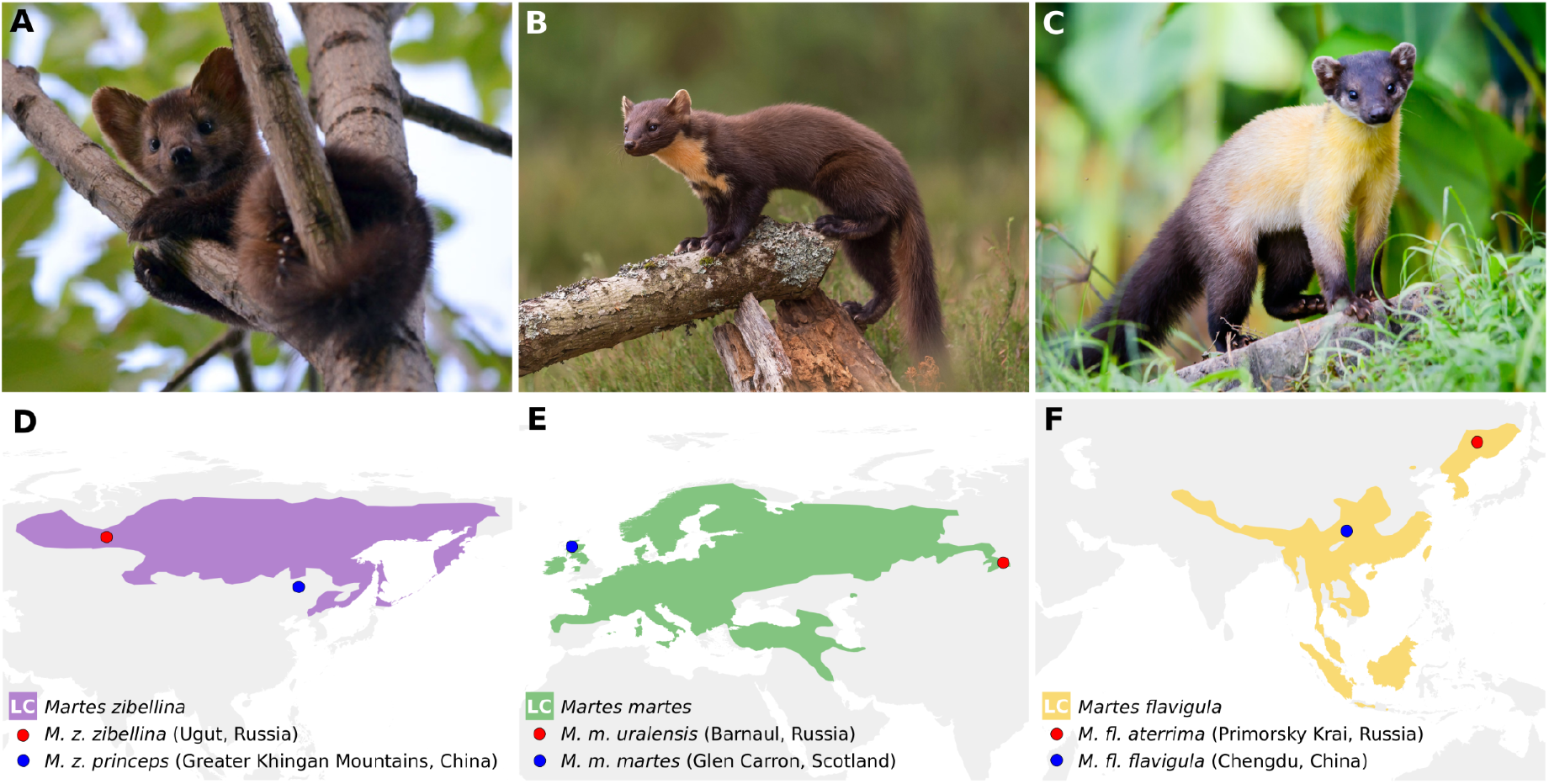
Pine marten (*M. martes)*, sable (*M. zibellina)*, yellow-throated marten (*M. flavigula*) and their ranges. A – Sable by E. Medvedeva (Wikimedia Commons, CC BY-SA 3.0); B – Pine marten by Caroline Legg (Flickr, CC BY 2.0); C – Yellow-throated marten by Rushen (Flickr, CC BY-SA 2.0); D-F – Ranges and global conservation statuses of three *Martes* species based on data from The International Union for Conservation of Nature’s Red List of Threatened Species (version 2025-1). Global conservation status: LC – Least Concern. Blue dots show the locations of individuals used for previously published genome assemblies, and red dots show the locations of individuals used for the genome assemblies reported in this study.

Resolving questions of taxonomy and intraspecific genetic structure within *Martes* species requires whole-genome sequencing, as has been repeatedly demonstrated in multiple studies of Mustelidae [26, 27]. Although genomic data for several *Martes* species have become available in recent years, the geographic distribution of the sequenced samples remains narrow and fails to capture the full extent of intraspecific variation. To date, genome assemblies have been published for a limited number of *Martes* species: a scaffold-level assembly of *M. z. princeps* (GCA_012583365.1, Greater Khingan Mountains, China) [28] and four chromosome-length assemblies for *M. f. toufoeus* (GCA_040938555.1, Gansu Province, China, short reads) [29], *M. m. martes* (GCA_963455335.1, Glen Carron, Scotland, long reads) [30], *M. f. foina* (GCA_964304585.1, Laze, Slovenia, long reads) [31], and *M. fl. flavigula* (GCA_029410595.1, Chengdu, Sichuan Province, China, long reads) [32]. These assemblies provide a valuable foundation but represent only single populations or subspecies and therefore do not allow a comprehensive reconstruction of intraspecific diversity.

In this study, we present chromosome-length assemblies for three *Martes* subspecies: *M. z. zibellina, M. m. uralensis*, and *M. fl. aterrima*. Unlike previously published genomes, these assemblies originate from subspecies inhabiting different and often more isolated parts of the species’ ranges (Figure **1**), including typical and peripheral forms (*M. uralensis* and *M. fl. aterrima*). By complementing existing resources, they broaden both the geographic and taxonomic coverage of genomic data for *Martes*.

## Materials and methods

### Samples and DNA extraction

To generate data for *de novo* assemblies, we used primary fibroblast cell lines from a female Tobol sable, *M. z. zibellina* (MZIB1f, 2019-0249, sample origin Uvat, Khanty-Mansi Autonomous Okrug – Yugra, Russia), a female pine marten from the Altai region, *M. m. uralensis* (MMAR1, 2018-0022, sample origin Barnaul Zoo, Russia) and a male Far Eastern yellow-throated marten from Primorsky Krai, *M. fl. aterrima* (MFLA2m, 2018-0732, sample origin Novosibirsk Zoo, Russia), which were obtained from the Novosibirsk Cell Line Collection located at the Institute of Molecular and Cellular Biology, Siberian Branch of the Russian Academy of Sciences (IMCB SB RAS). The origin of the zoo animals was confirmed by staff of both zoos. Sample collection, transportation and cell line establishment were previously described in detail [33]. DNA extraction was performed using the standard phenol-chloroform protocol [34]. For *de novo* assembly of all three genomes, we generated two types of libraries: linked reads and Hi-C. The linked read libraries were prepared using the Chromium Genome Reagent Kit version2 and the microfluidic Genome Chip run in a Chromium Controller instrument according to the manufacturer’s instructions (10X Genomics, Pleasanton, California, USA). The Hi-C libraries were prepared according to the original protocol [35]. All prepared libraries were sequenced with paired-end 150 bp reads on the Illumina NovaSeq 6000 or Illumina HiSeq X Ten platforms. All manipulations with the samples were performed according to the permission of IMCB Ethical Committee № 01/21 issued on 26 January 2021.

### De novo genome assembly

*De novo* assembly of each of three genomes was performed in four stages. First, we generated draft assemblies from the linked-read Illumina sequencing data using the Supernova v2 [36] assembler. Next, we scaffolded assemblies to chromosome-level using the Hi-C sequencing data with Juicer v1.6 [37] and 3D-DNA v180419 [38] with the default parameters. For the third step we manually curated the assemblies in Juicebox v2.16.00 [37] to correct misjoins. Finally, haplotype duplications were detected using purge_dups v1.2.6 [39], based on sequence similarity and coverage. To avoid overpurging, we removed duplicates located only on non-chromosomal scaffolds. In cases when all the copies were located in non-chromosomal scaffolds, the longest one was retained. Completeness of the genome assemblies were assessed with BUSCO v5.4.2 [40] using the database Mammalia_odb v10, 2021-02-19.

### Repeats, whole-genome alignment, chromosomes and inversions

Nomenclature of chromosomal scaffolds in the genome assemblies of the *M. z. zibellina* (this study), *M. m. uralensis* (this study), *M. m. martes*, GCA_963455335.1 [30], *M. f. foina*, GCA_964304585.1 [31], *M. fl. atterima* (this study) and *M. fl. flavigula*, GCA_029410595.1 [32], was defined via a comparison of the whole-genome alignment (WGA) that included the genome assemblies of *M. f. toufoeus* (GCA_040938555.1) [29] and chromosome painting maps [41, 42]. First, tandem and dispersed repeats in the genome assemblies were identified using Tandem Repeats Finder v4.09.1 [43] with parameters “2 7 7 80 10 50 2000 -l 10”, WindowMasker v1.0.0 [44] with default parameters, and RepeatMasker v4.1.2.p1 [45] with the parameter “-species carnivora”. Tandem Repeats Finder and RepeatMasker were run using the Dfam TETools v1.88.5 container (https://github.com/Dfam-consortium/TETools). Subsequently, BEDTools v2.31.0 [46] with the “-soft” parameter was employed to softmask the genome assemblies using the identified repeat elements. Then, we performed a multiple whole-genome alignment of these masked genome assemblies using Progressive Cactus v2.8.0 [47] with default parameters. Next, we extracted synteny blocks from the multiple alignment using halSynteny v2.2 [48] with the options “--minBlockSize 50000 --maxAnchorDistance 50000”, visualized and categorized (translocated, inverted or “normal”) the obtained synteny blocks using ChromoDoter v0.4 [49] and the scripts (*draw_synteny*.*py* and *draw_macrosynteny*.*py*) from the MACE v1.1.32 package [50], and assigned the chromosome names to the scaffolds. We also transferred coordinates of centromeres from the *M. f. toufoeus* assembly, for which approximate locations were previously reported [29], as the chromosomes of *M. zibellina, M. martes* and *M. flavigula* have similar positions of centromeres [41, 42, 51]. Finally, we compared G-banding [41, 42] of all four *Martes* species to verify candidate inversions on chr11, chr12/chr15 and chr18.

### Prediction of protein-coding genes

We predicted protein-coding genes in the assemblies of *M. z. zibellina, M. m. uralensis*, and *M. fl. aterrima* using the BRAKER v3.0.8 [52] pipeline. For the annotations of *M. z. zibellina* and *M. m. uralensis*, we used previously generated RNA-seq data from both these species (Supplementary Table ST2) [28, 30, 53]. For *M. fl. aterrima*, only RNA-seq data of this species were used (Supplementary Table ST2) [32]. Additional inputs included the BUSCO v5.4.2 [40] Mammalia_odb10 database (2024-01-08), and protein hints from the Metazoa database of OrthoDB v11, generated using the orthodb-clades pipeline (https://github.com/tomasbruna/orthodb-clades) [54]. Gene prediction was performed using GeneMark-ETP v1.02 [55], which was trained on RNA-seq and protein homology data, while AUGUSTUS v3.5.0 [56, 57] provided further gene prediction supported by the external data. To assign gene names to the predicted gene models (i.e., functional annotation), we used eggNOG-mapper v2.1.12 and the EggNOG v5.0 database (Mammalia subset) [58, 59]. Versions of all used tools and databases used for raw read data processing, genome assembly, synteny analysis, and annotation are listed in Table 1.

**Table 1.** Tools used for assembly, annotation, and analysis of the genomes.

| Stage of analysis | Software/Database | Version | Reference |
| --- | --- | --- | --- |
| Data QC | KrATER | v2.5 | https://github.com/mahajrod/krater |
|  | Jellyfish | v2.2.10 | [60] |
|  | GenomeScope2 | v2.0 | [61] |
| Genome assembly | Supernova | v2 | [36] |
|  | Juicer | v2019 | [62] |
|  | 3D-DNA | v2019 | [38] |
|  | Juicebox Assembly Tools | v2019 | [37] |
| Genome assembly QC | BUSCO | v5.5.0 | [40] |
|  | OrthoDB* | odb10 | [63] |
| Repeat detection and masking | RepeatMasker | v4.1.6 | [45] |
|  | Dfam* | v3.8 | [64] |
|  | Windowmasker | v1.0.0 | [44] |
|  | TRF | v4.09.1 | [43] |
|  | Bedtools | v2.29 | [46] |
| Whole genome alignment | LAST | v981 | [65] |
|  | ProgressiveCactus | v1.0 | [47] |
|  | halSynteny | v2.2 | [48] |
| Gene prediction | BRAKER | v3.0.8 | [52] |
|  | AUGUSTUS | v3.5.0 | [56, 57] |
|  | eggNOG-mapper | v2.1.12 | [58] |
|  | eggNOG (Mammalia)* | v5.0 | [59] |
| Visualization | MAVR | v0.113 | https://github.com/mahajrod/mavr |
|  | MACE | v1.1.32 | https://github.com/mahajrod/mace |

\* databases

## Results and discussion

### Chromosome-length genome assemblies

We generated and assembled 619,134,368 linked reads (49.7x coverage) and 605,828,662 Hi-C reads from a female Tobol sable (MZIB, *M. z. zibellina*), 633,013,226 linked reads (49.6x) and 687,528,994 Hi-C reads from a female Ural pine marten (MMAR, *M. m. uralensis*), and 637,842,932 linked reads (48.4x) and 832,218,710 Hi-C reads from a male Far Eastern yellow-throated marten (MFLA, *M. fl. aterrima*). The resulting chromosomal-length reference assemblies have total lengths of 2.39 Gbp, 2.40 Gbp, and 2.45 Gbp for the *M. z. zibellina, M. m. uralensis* and *M. fl. aterrima*, respectively, which closely match the 23-mer based estimates (2.4 Gbp, 2.45 Gbp and 2.46 Gbp, Supplementary Figure SF1). All three genomes exhibit identical GC contents (41.3 %). The scaffold N50 values for the *M. z. zibellina, M. m. uralensis* and *M. fl. aterrima* assemblies are 143.6 Mbp, 144.6 Mbp and 137.4 Mbp (Supplementary Table ST3), respectively, reflecting the lengths of individual chromosomes. Each assembly comprises a number of chromosomal scaffolds corresponding to the chromosome pairs in each species’ karyotype (2n = 38 for *M. z. zibellina* and *M. m. uralensis*; 2n = 40 for *M. fl. aterrima*), affirming the chromosome-length status of all assemblies. BUSCO analysis (Supplementary Table ST4, Mammalia_odb v.10 with 9,226 BUSCOs) demonstrated high completeness for *M. z. zibellina* (96.1% complete BUSCOs) and *M. m. uralensis* (96.4 %) assemblies. For *M. zibellina*, it is a significant improvement compared to a previously published assembly [28] of a sample representing the subspecies *M. z. princeps* (94.8% complete BUSCOs), which is also highly fragmented and notably below chromosome-level (scaffold N50 5.2 Mbp), whereas most parameters of our *M. m. uralensis* assembly are similar to the available genome [30] of *M. m. martes* from Scotland (scaffold N50 146.29, 96.3 % complete BUSCOs). However, in our *M. fl. aterrima* assembly we found a lower fraction of the complete BUSCOs (93.6 %) than in the published [32] *M. fl. flavigula* genome (96.9 %).

### Macrosynteny

The recently published stone marten, *M. f. toufoeus* (MFOI), genome assembly [29] and comparative chromosome painting maps of all four marten species [41, 42] allowed us to connect the assemblies with each species’ karyotype. We detected no discrepancies between the cytogenetic data and whole-genome alignments. For each chromosome of all subspecies (except *M. z. princeps* due to the fragmented assembly), we identified the corresponding chromosomal scaffold in the assembly (Supplementary Table ST5). Whole-genome alignment showed that multiple chromosomal scaffolds in our assemblies have different orientations (Figure 2D).

**Figure 2.**
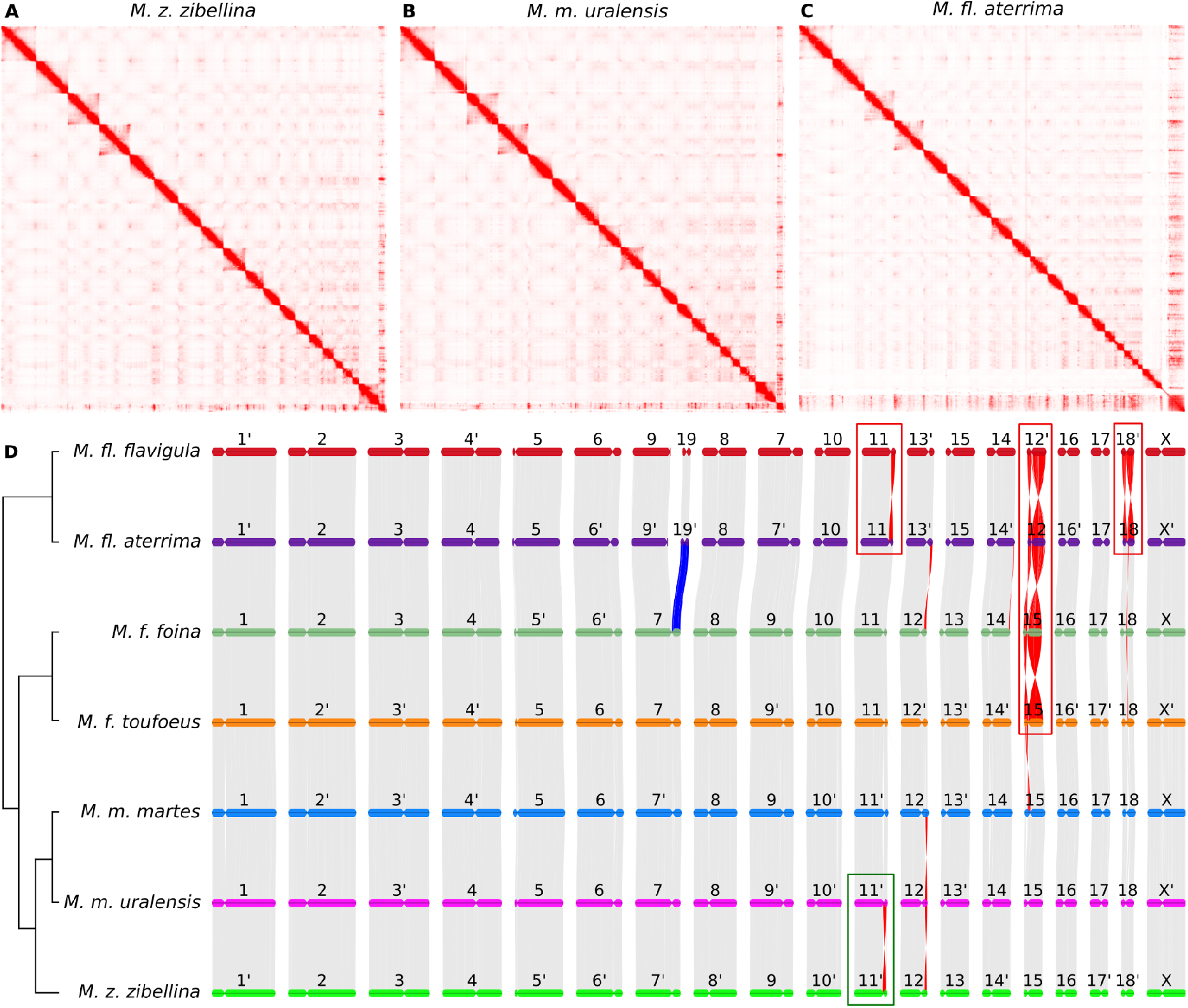
Hi-C maps and macrosynteny between marten species and subspecies. A, B, C – Hi-C contact maps of *M. z. zibellina* (Tobol sable), *M. m. uralensis* (Ural pine marten) and *M. fl. aterrima (*Far East yellow-throated marten). Autosomes are arranged according to length from the top left to bottom right, followed by the chrX; D – macro-level synteny map between seven subspecies of four marten species. Colored horizontal blocks represent individual chromosomes. Vertical gray lines represent non-inverted syntenic blocks. Large inversions (>1 Mbp) are highlighted in red. Chromosomes labeled by primes (‘) were reverse complemented to follow the orientation of corresponding homologs in the *M. foina* assembly. Green rectangle indicates the cytogenetically verified inversion on chr 11, red rectangles indicate confirmed missassemblies. The cladogram of the species [68, 69] is shown to the left. Note that the nomenclature of the *M. fl. ssp* chromosomes are slightly different from the other marten species.

Among the species, we identified two large-scale rearrangements of the same type (Figure 2D, red rectangles) on chr18 and chr15 (chr12 in *M. fl. ssp*). However, such a pattern (a simultaneous inversion of both chromosomal arms or a telomeric join) is a common artifact in assemblies, when the Hi-C signal over centromere (or other large repetitive region missing in the assembly) is weak [66, 38], and sometimes it is difficult to detect even during an intensive manual curation. Given the distribution of the putative artifacts among the (sub)species and that the *M. f. foina* and *M. fl. flavigula* assemblies are Nanopore-based, we assumed that misassemblies are present in the two latter genomes, as insufficient polishing of contigs can result in a lower mapping rate [67] of the Hi-C reads, and, in turn, in the reduced Hi-C signal during scaffolding. Among the five other (sub)species assemblies, four are short-read based, which also can lead to large-scale artifacts due to higher fragmentation of the contigs. However, we observed no such large-scale rearrangements between these assemblies and the HiFi-based *M. m. martes* assembly (Figure 2D). Comparing published G-banded karyotypes of all four species [42, 41], we were not able to prove or disprove these rearrangements. The small size of the regions affected by rearrangements and low number of G-bands in the investigated areas does not allow for a definitive validation of the inversions on chr18 and chr15 (chr12 in *M. fl. ssp*) (Supplementary Figure SF2). However, we reconstructed Hi-C contact maps for *M. f. foina* and *M. fl. flavigula* using the original data used for assembly [31, 32] and found that all these putative missassemblies are indeed artefacts of the Hi-C scaffolding (Supplementary Figures SF3-SF7).

We confirmed a sable-specific inversion on chr11 (11.5 Mbp, Figure 2D, green rectangle) via comparison of published karyotypes [41] that is accompanied by a change in centromeric position (acrocentric in *M. z. zibellina* and subtelocentric in all other *Martes* species). We found a similar inversion between the assemblies of *M. fl. ssp* (Figure 2D, red rectangle), but G-banded chr11 of *M. fl. aterrima* [41] (the same individual was used to generate the assembly) and *M. fl. flavigula* [42] have the same chromosome morphology (subtelocentric) and a similar number and distribution of G-bands, which is closer to *M. m. uralensis* and *M. f. toufoeus* than to *M. z. zibellina* (Supplementary Figure SF2). The reconstructed Hi-C map of *M. fl. flavigula* (Supplementary figure SF2) confirmed that it is an artefact (an inversion of the p-arm) of this assembly. The remaining inversions were impossible to check because they were all too small for cytogenetic verification, but we found no contradictions with Hi-C contact maps.

### Repeats, pseudoautosomal region and protein-coding genes

We detected comparable proportions of transposable elements across all newly generated and publicly available *Martes* genome assemblies (Supplementary Table ST6). The overall fraction of interspersed repeats ranged from 36.9% in *M. z. princeps* to 40.4% in *M. fl. flavigula*, with the majority of repetitive content contributed by retroelements, primarily SINEs and LINEs. Within the sables, the genome assembly of the *M. z. zibellina* harbored 39.3% interspersed repeats, which is slightly higher than in the publicly available *M. z. princeps* assembly (36.9%), mainly due to LINEs (21.7% vs. 19.9%) and SINEs (9.9% vs. 9.4%). For the yellow-throated martens, both subspecies showed the highest levels of repetitive content. The *M. fl. aterrima* assembly contained 39.7% interspersed repeats, while *M. fl. flavigula* harbored 40.4%, due to a slightly higher fraction of LINEs (22.1% vs. 22.7%) in the latter.

Kimura divergence profiles confirmed these observations (Figure **3**). The profiles of *M. f. toufoeus* and *M. f. foina* were practically indistinguishable, while for other species the differences were mainly in LINE abundance. The publicly available assemblies of *M. martes* and *M. flavigula* contained slightly more LINEs, whereas for *M. zibellina*, the our chromosome-level and linked read-based genome assembly showed a higher fraction than the published scaffold-level and mate pair-based assembly [28], which underrepresents repeats. Taken together, these results indicate that the composition of interspersed repeats in *Martes* is highly conserved across species and subspecies, with the total amount consistently accounting for 37–40% of the genome, and LINEs (~20–23%) and SINEs (~9–10%) representing the predominant classes, which is typical for carnivorans.

**Figure 3.**
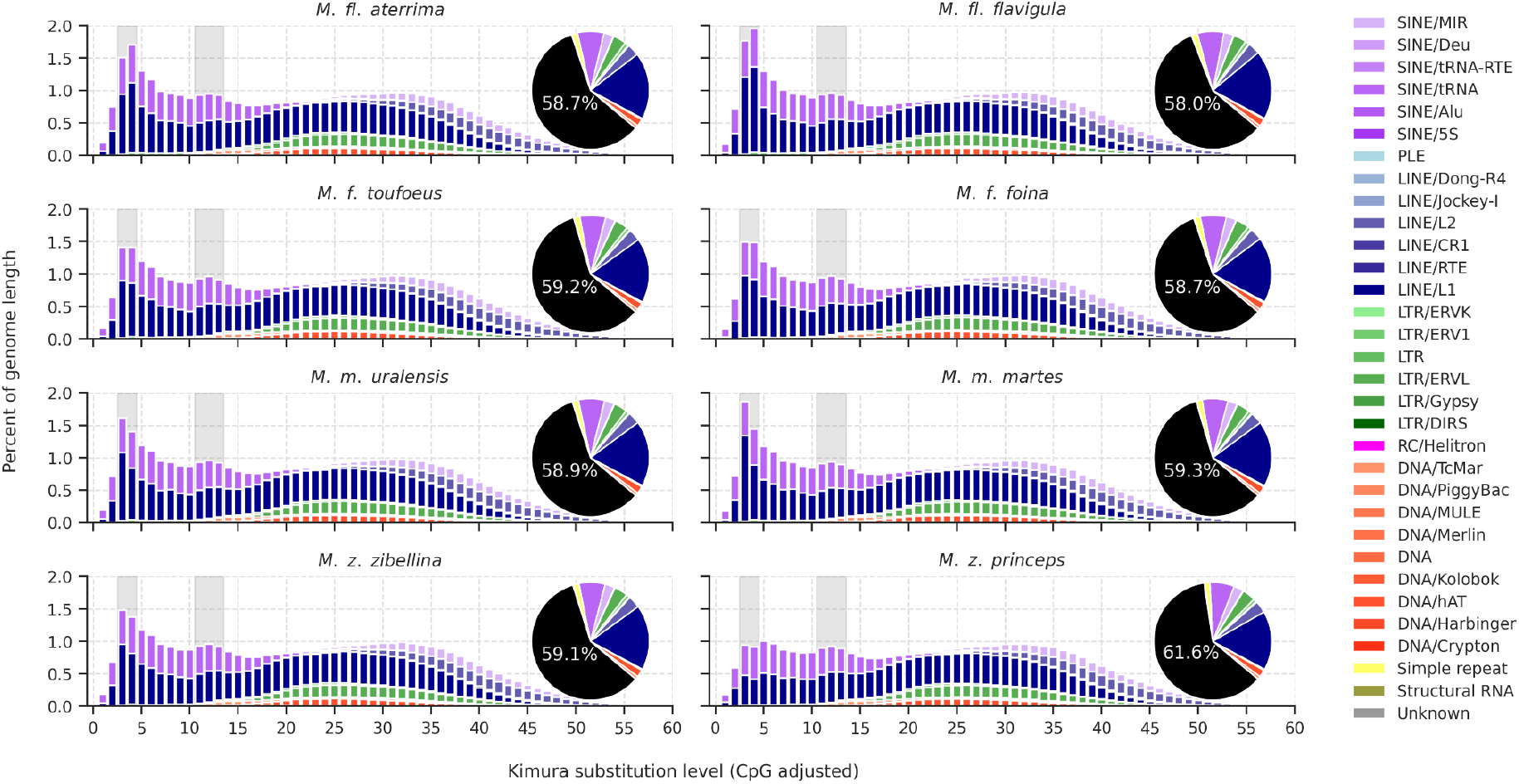
Kimura divergence profiles of transposable elements in eight genome assemblies of *Martes* subspecies. Profiles of the three newly-generated assemblies are shown on the left, and publicly available assemblies on the right. The graphs display the distribution and divergence of major transposable elements classes. Pie charts show the proportions of different repeat element classes, and for the non-repetitive fraction (black), the genome percentage is indicated.

We identified the coordinates of the pseudoautosomal region (PAR) in the genome assemblies generated in this study. In all assemblies, the PAR, as expected, was located at the end of the X chromosome. However, for *M. fl. aterrima*, it has a length of 5.02 Mbp (HiC_scaffold_20:116730000-121750000), which is notably smaller than the estimated values for the PAR in *M. z. zibellina* and the *M. m. uralensis* (6.48 Mbp and 6.45 Mbp, respectively). However, in this case, these differences in PAR size do not reflect interspecific variation but rather stem from the difficulty of accurately defining its boundaries due to uneven coverage (Supplementary Figure SF8).

We predicted 20,158 protein-coding genes in the *M. z. zibellina* assembly, 20,259 in the *M. m. uralensis* assembly, and 20,521 in the *M. fl. aterrima* assembly. Of these, over 90% were functionally annotated, with 18,300 named genes in the *M. z. zibellina*, 18,248 in the *M. m. uralensis*, and the 18,595 in *M. fl. aterrima* (Supplementary File SF1). In total, 14,978 named genes were shared between all three assemblies (Supplementary Figure SF9). BUSCO analysis indicated high gene set completeness, with scores of 97.7% for *M. z. zibellina* and *M. m. uralensis*, and 95.1% for *M. fl. aterrima*, confirming the overall high quality of the predictions (Supplementary Table ST7).

## Conclusions

We generated and annotated chromosome-length assemblies for three species within the genus *Martes*, covering new subspecies from distant or isolated parts of the respective ranges of each species. For example, the sampled individuals of *M. zibellina* used to generate our assembly (from Khanty-Mansi Autonomous Okrug– Yugra, Russia) and the previously published assembly (from the Greater Khingan Mountains, China [28]) originated from regions separated by ~4’000 km (Figure 1D), whereas the samples used for the two *M. martes* assemblies (from Barnaul, Russia versus Glen Carron, Scotland, United Kingdom [30]) are separated by a distance of ~6’000 km (Figure 1E). For the two *M. flavigula* assemblies, the geographical distance is smaller (~2’700 km), but the corresponding parts of the range are highly isolated (Figure 1F), and there is an ongoing debate within the scientific community whether *M. fl. flavigula* and the *M. fl. aterrima* should be treated as distinct species [24]. Therefore, the increase of available marten genomes representing new subspecies provides a foundation for more accurate assessments of intraspecific variation (via mapping to closer related references) and opens the possibility to reinvestigate the taxonomy of the genus on the whole-genome level.

Analyses of the synteny between our and previously published genome assemblies identified several large-scale misassemblies in the previously published Nanopore-based assemblies of *M. fl. flavigula* (chr11, chr12, chr18) and *M. f. foina* (chr15) [31, 32]. Our results highlight that extensive curation is still necessary even for long-read based assemblies and that a comparative approach based on whole genome alignment should be a mandatory part of it. Synteny-based analyses help to reveal specific patterns of assembly artefacts, for example, telomeric joints and inverted chromosomal arms, and highlights putative misassemblies for further investigation. Finally, by adding the published cytogenetic data associated with each marten species, we identified a highly confident inversion between the *M. zibellina* and *M. martes*, which encompasses the whole p-arm of chr11. Given the known hybridization between these species and putative fertility issues in hybrids [70, 71], the occurrence and frequency of this rearrangement requires further investigation using a wider and larger sampling at the geographical and population level.

## Supplementary

SupplementaryFiguresAndTables: link

SupplementaryFiles: link

## Acknowledgements

We thank Sergei Pisarev and Pavel Reznichenko from the Barnaul Zoo “Lesnayia skazka” (eng. “Forest tale”, Barnaul, Russia) for providing samples used to generate the *Martes martes uralensis* cell line. We acknowledge team of Rostislav Shilo Novosibirsk Zoo (Russia, Novosibirsk), in particular Olga Shilo (deputy director), Rosa Solovyova (head of carnivore department) and Svetlana Verkholantseva (veterinarian), for for providing samples used to generate the *Martes flavigula aterrima* cell line. The research reported in this study was partially completed using equipment (materials) belonging to the large-scale research facilities of the “Cryobank of cell cultures” at the Institute of Molecular and Cellular Biology SB RAS (Novosibirsk, Russia). We are especially grateful to M. Thomas P. Gilbert for his brilliant advice and generous help.

## Data availability

The linked reads used for *de novo* assemblies are available from BioProject PRJNA905543. The Hi-C data is available under accessions SRR16970334, SRR16086878 and SRR16086880, for *Martes zibellina zibellina, Martes martes uralensi*s and *Martes flavigula aterrima*, respectively. Assemblies are available from the NCBI Genome database.

## Literature

1. Wozencraft WC. Order Carnivora. In: Wilson DE, Reeder DM, editors. Mammal Species of the World: a Taxonomic and Geographic Reference. 3rd edition. Baltimore: The Johns Hopkins University Press; 2005. p. 532–628.

2. Koepfli K-P, Deere KA, Slater GJ, Begg C, Begg K, Grassman L, et al. Multigene phylogeny of the Mustelidae: Resolving relationships, tempo and biogeographic history of a mammalian adaptive radiation. BMC Biol. 2008;6:10. 10.1186/1741-7007-6-10.

3. Sato JJ, Wolsan M, Prevosti FJ, D’Elía G, Begg C, Begg K, et al. Evolutionary and biogeographic history of weasel-like carnivorans (Musteloidea). Mol Phylogenet Evol. 2012;63:745–57. 10.1016/j.ympev.2012.02.025.

4. Monakhov VG. IUCN Red List of Threatened Species: Martes zibellina. IUCN Red List Threat Species. 2015. 10.2305/IUCN.UK.2016-1.RLTS.T41652A45213477.en.

5. Herrero J, Kranz A, Skumatov D, Abramov AV, Maran T, Monakhov VG. IUCN Red List of Threatened Species: Martes martes. IUCN Red List Threat Species. 2015. 10.2305/IUCN.UK.2016-1.RLTS.T12848A45199169.en.

6. Abramov AV, Kranz A, Herrero J, Choudhury AU, Maran T. IUCN Red List of Threatened Species: Martes foina. IUCN Red List Threat Species. 2015. 10.2305/IUCN.UK.2024-2.RLTS.T29672A259348828.en.

7. Abramov AV, Kaneko Y, Masuda R. IUCN Red List of Threatened Species: Martes melampus. IUCN Red List Threat Species. 2015. 10.2305/IUCN.UK.2015-4.RLTS.T41650A45213228.en.

8. Helgen K, Reid F. IUCN Red List of Threatened Species: Martes americana. IUCN Red List Threat Species. 2015. 10.2305/IUCN.UK.2016-1.RLTS.T41648A45212861.en.

9. Colella JP, Lan T, Talbot SL, Lindqvist C, Cook JA. Whole-genome resequencing reveals persistence of forest-associated mammals in Late Pleistocene refugia along North America’s North Pacific Coast. J Biogeogr. 2021;48:1153–69. 10.1111/jbi.14068.

10. Chutipong W, Duckworth JW, Timmins RJ, Choudhury AU, Abramov AV, Roberton S, et al. IUCN Red List of Threatened Species: Martes flavigula. IUCN Red List Threat Species. 2015. 10.2305/IUCN.UK.2024-2.RLTS.T41649A259348347.en.

11. Mudappa D, Jathana D, Raman TRS. IUCN Red List of Threatened Species: Martes gwatkinsii. IUCN Red List Threat Species. 2015. 10.2305/IUCN.UK.2015-4.RLTS.T12847A45199025.en.

12. Monakhov VG. Martes zibellina (Carnivora: Mustelidae). Mamm Species. 2011;43:75–86. 10.1644/876.1.

13. Pocock RI. The Oriental Yellow-throated Marten (Lamprogale). By RI POCOCK, FRS, FZS. In: Proceedings of the Zoological Society of London. Wiley Online Library; 1936. p. 531–53.

14. Yudin VG, Yudina EV. The Yellow-Throated Marten of the Far East of Russia. Federal Scientic Center of the East Asia Terrestrial Biodiversity, Far Eastern Branch, RAS; 2022.

15. Monakhov VG. Martes martes (Carnivora: Mustelidae). Mamm Species. 2022;54:seac007. 10.1093/mspecies/seac007.

16. Mammal Diversity Database. 2025. 10.5281/zenodo.17033774.

17. Kamada S, Moteki S, Baba M, Ochiai K, Masuda R. Genetic distinctness and variation in the Tsushima Islands population of the Japanese marten, Martes melampus (Carnivora: Mustelidae), revealed by microsatellite analysis. Zoolog Sci. 2012;29:827–33.

18. Jo Y-S, Baccus JT, Koprowski JL. Mammals of Korea. National Institute of Biological Resources; 2018.

19. Tsoupas A, Andreadou M, Papakosta MA, Karaiskou N, Bakaloudis DE, Chatzinikos E, et al. Phylogeography of Martes foina in Greece. Mamm Biol. 2019;95:59–68. 10.1016/j.mambio.2019.02.004.

20. Ranyuk M, Modorov M, Monakhov V, Devyatkin G. Genetic differentiation of autochthonous sable populations in Western and Eastern Siberia. J Zool Syst Evol Res. 2021;59:2539–52. 10.1111/jzs.12565.

21. Li B, Lu J, Monakhov V, Kang H, Xu Y, An B, et al. Phylogeography of subspecies of the sable (Martes zibellina L.) based on mitochondrial genomes: implications for evolutionary history. Mamm Biol. 2021;101:105–20. 10.1007/s42991-020-00092-0.

22. Filimonov PA, Zakharov ES, Kochkarev PV, Politov DV, Davydov AV, Onokhov AA, et al. Genetic structure of native and naturalized populations of sable (Martes zibellina L.) of the Central Siberian Plateau and adjacent territories. Biol Invasions. 2024;26:2139–51. 10.1007/s10530-024-03299-1.

23. Dawson NG, Colella JP, Small MP, Stone KD, Talbot SL, Cook JA. Historical biogeography sets the foundation for contemporary conservation of martens (genus Martes) in northwestern North America. J Mammal. 2017;98:715–30. 10.1093/jmammal/gyx047.

24. Matyuschkin E.N. Yellow-throated marten (Martes (Charronia) flavigula Boddaert, 1785), (Mustelidae, Carnivora) in the Russian Far East. Lutreola Mosc. 1993;:2–9.

25. Rozhnov VV. Taxonomic notes on the yellow-throated marten Martes flavigula. Zool J. 1995;74:131–8.

26. Colella JP, Frederick LM, Talbot SL, Cook JA. Extrinsically reinforced hybrid speciation within Holarctic ermine (Mustela spp.) produces an insular endemic. Divers Distrib. 2021;27:747–62.

27. de Ferran V, Figueiró HV, de Jesus Trindade F, Smith O, Sinding M-HS, Trinca CS, et al. Phylogenomics of the world’s otters. Curr Biol. 2022;32:3650–3658.e4. 10.1016/j.cub.2022.06.036.

28. Liu G, Zhao C, Xu D, Zhang H, Monakhov V, Shang S, et al. First Draft Genome of the Sable, Martes zibellina. Genome Biol Evol. 2020;12:59–65. 10.1093/gbe/evaa029.

29. Tomarovsky A, Khan R, Dudchenko O, Totikov A, Serdyukova NA, Weisz D, et al. Chromosome-length genome assembly of the stone marten (Martes foina, Mustelidae): A new view on one of the cornerstones in carnivore cytogenetics. J Hered. 2025;116:548–57. 10.1093/jhered/esaf001.

30. O’Brien D, Januszczak I, Natural History Museum Genome Acquisition Lab, Darwin Tree of Life Barcoding collective, Wellcome Sanger Institute Tree of Life Management, Samples and Laboratory team, Wellcome Sanger Institute Scientific Operations: Sequencing Operations, et al. The genome sequence of the European pine marten, Martes martes (Linnaeus, 1758). Wellcome Open Res. 2024;9:325. 10.12688/wellcomeopenres.22458.1.

31. Martes foina genome assembly mMarFoi2.1. NCBI. https://www.ncbi.nlm.nih.gov/datasets/genome/GCA_964304585.1/. Accessed 4 Sept 2025.

32. Mei X, Liu G, Yan J, Zhao C, Wang X, Zhou S, et al. A chromosome-level genome assembly of the yellow-throated marten (Martes flavigula). Sci Data. 2023;10:216. 10.1038/s41597-023-02120-3.

33. Beklemisheva VR, Perelman PL, Lemskaya NA, Kulemzina AI, Proskuryakova AA, Burkanov VN, et al. The Ancestral Carnivore Karyotype As Substantiated by Comparative Chromosome Painting of Three Pinnipeds, the Walrus, the Steller Sea Lion and the Baikal Seal (Pinnipedia, Carnivora). PLOS ONE. 2016;11:e0147647. 10.1371/journal.pone.0147647.

34. Sambrook J, Russell DW. Purification of nucleic acids by extraction with phenol: chloroform. Cold Spring Harb Protoc. 2006;2006:pdb.prot4455.

35. Rao SSP, Huntley MH, Durand NC, Stamenova EK, Bochkov ID, Robinson JT, et al. A 3D Map of the Human Genome at Kilobase Resolution Reveals Principles of Chromatin Looping. Cell. 2014;159:1665–80. 10.1016/j.cell.2014.11.021.

36. Weisenfeld N, Kumar V, Shah P, Church D, Jaffe D. Direct determination of diploid genome sequences. Genome Res. 2017;27. 10.1101/gr.214874.116.

37. Dudchenko O, Shamim MS, Batra SS, Durand NC, Musial NT, Mostofa R, et al. The Juicebox Assembly Tools module facilitates de novo assembly of mammalian genomes with chromosome-length scaffolds for under $1000. bioRxiv. 2018;:254797. 10.1101/254797.

38. Dudchenko O, Batra SS, Omer AD, Nyquist SK, Hoeger M, Durand NC, et al. De novo assembly of the Aedes aegypti genome using Hi-C yields chromosome-length scaffolds. Science. 2017;356:92–5. 10.1126/science.aal3327.

39. Guan D, McCarthy SA, Wood J, Howe K, Wang Y, Durbin R. Identifying and removing haplotypic duplication in primary genome assemblies. Bioinformatics. 2020;36:2896–8. 10.1093/bioinformatics/btaa025.

40. Manni M, Berkeley MR, Seppey M, Zdobnov EM. BUSCO: Assessing Genomic Data Quality and Beyond. Curr Protoc. 2021;1:e323. 10.1002/cpz1.323.

41. Beklemisheva VR, Lemskaya NA, Prokopov DY, Perelman PL, Romanenko SA, Proskuryakova AA, et al. Maps of Constitutive-Heterochromatin Distribution for Four Martes Species (Mustelidae, Carnivora, Mammalia) Show the Formative Role of Macrosatellite Repeats in Interspecific Variation of Chromosome Structure. Genes. 2023;14:489. 10.3390/genes14020489.

42. Nie W, Wang J, O’Brien PCM, Fu B, Ying T, Ferguson-Smith MA, et al. The genome phylogeny of domestic cat, red panda and five mustelid species revealed by comparative chromosome painting and G-banding. Chromosome Res. 2002;10:209–22. 10.1023/A:1015292005631.

43. Benson G. Tandem repeats finder: a program to analyze DNA sequences. Nucleic Acids Res. 1999;27:573–80. 10.1093/nar/27.2.573.

44. Morgulis A, Gertz EM, Schäffer AA, Agarwala R. WindowMasker: window-based masker for sequenced genomes. Bioinforma Oxf Engl. 2006;22:134–41. 10.1093/bioinformatics/bti774.

45. Tarailo-Graovac M, Chen N. Using RepeatMasker to Identify Repetitive Elements in Genomic Sequences. Curr Protoc Bioinforma. 2009;25:4.10.1–4.10.14. 10.1002/0471250953.bi0410s25.

46. Quinlan AR, Hall IM. BEDTools: a flexible suite of utilities for comparing genomic features. Bioinformatics. 2010;26:841–2. 10.1093/bioinformatics/btq033.

47. Armstrong J, Hickey G, Diekhans M, Fiddes IT, Novak AM, Deran A, et al. Progressive Cactus is a multiple-genome aligner for the thousand-genome era. Nature. 2020;587:246–51. 10.1038/s41586-020-2871-y.

48. Krasheninnikova K, Diekhans M, Armstrong J, Dievskii A, Paten B, O’Brien S. halSynteny: a fast, easy-to-use conserved synteny block construction method for multiple whole-genome alignments. GigaScience. 2020;9:giaa047. 10.1093/gigascience/giaa047.

49. Kliver S. ChromoDoter. 2022.

50. Kliver S. MACE v1.1.32. 2024.

51. Graphodatsky A, Perelman P, O’Brien SJ. Atlas of Mammalian Chromosomes. John Wiley & Sons, Incorporated; 2020.

52. Brůna T, Gabriel L, Hoff KJ. Navigating Eukaryotic Genome Annotation Pipelines: A Route Map to BRAKER, Galba, and TSEBRA. 2024. 10.48550/arXiv.2403.19416.

53. Xia T, Zhang L, Sun G, Yang X, Zhao C, Zhang H. Insights into cold tolerance in sable (Martes zibellina) from the adaptive evolution of lipid metabolism. Mamm Biol. 2021;101:861–70. 10.1007/s42991-021-00135-0.

54. Kuznetsov D, Tegenfeldt F, Manni M, Seppey M, Berkeley M, Kriventseva EV, et al. OrthoDB v11: annotation of orthologs in the widest sampling of organismal diversity. Nucleic Acids Res. 2023;51:D445–51. 10.1093/nar/gkac998.

55. Brůna T, Lomsadze A, Borodovsky M. GeneMark-ETP significantly improves the accuracy of automatic annotation of large eukaryotic genomes. Genome Res. 2024;34:757–68. 10.1101/gr.278373.123.

56. Stanke M, Keller O, Gunduz I, Hayes A, Waack S, Morgenstern B. AUGUSTUS: ab initio prediction of alternative transcripts. Nucleic Acids Res. 2006;34 suppl_2:W435–9. 10.1093/nar/gkl200.

57. Stanke M, Diekhans M, Baertsch R, Haussler D. Using native and syntenically mapped cDNA alignments to improve de novo gene finding. Bioinformatics. 2008;24:637–44. 10.1093/bioinformatics/btn013.

58. Cantalapiedra CP, Hernández-Plaza A, Letunic I, Bork P, Huerta-Cepas J. eggNOG-mapper v2: Functional Annotation, Orthology Assignments, and Domain Prediction at the Metagenomic Scale. Mol Biol Evol. 2021;38:5825–9. 10.1093/molbev/msab293.

59. Huerta-Cepas J, Szklarczyk D, Heller D, Hernández-Plaza A, Forslund SK, Cook H, et al. eggNOG 5.0: a hierarchical, functionally and phylogenetically annotated orthology resource based on 5090 organisms and 2502 viruses. Nucleic Acids Res. 2019;47:D309–14. 10.1093/nar/gky1085.

60. Marçais G, Kingsford C. A fast, lock-free approach for efficient parallel counting of occurrences of k-mers. Bioinformatics. 2011;27:764–70. 10.1093/bioinformatics/btr011.

61. Ranallo-Benavidez TR, Jaron KS, Schatz MC. GenomeScope 2.0 and Smudgeplot for reference-free profiling of polyploid genomes. Nat Commun. 2020;11:1432. 10.1038/s41467-020-14998-3.

62. Durand NC, Shamim MS, Machol I, Rao SSP, Huntley MH, Lander ES, et al. Juicer Provides a One-Click System for Analyzing Loop-Resolution Hi-C Experiments. Cell Syst. 2016;3:95–8. 10.1016/j.cels.2016.07.002.

63. Kriventseva EV, Kuznetsov D, Tegenfeldt F, Manni M, Dias R, Simão FA, et al. OrthoDB v10: sampling the diversity of animal, plant, fungal, protist, bacterial and viral genomes for evolutionary and functional annotations of orthologs. Nucleic Acids Res. 2019;47:D807–11. 10.1093/nar/gky1053.

64. Storer J, Hubley R, Rosen J, Wheeler TJ, Smit AF. The Dfam community resource of transposable element families, sequence models, and genome annotations. Mob DNA. 2021;12:2. 10.1186/s13100-020-00230-y.

65. Frith MC, Kawaguchi R. Split-alignment of genomes finds orthologies more accurately. Genome Biol. 2015;16:1–17.

66. Burton JN, Adey A, Patwardhan RP, Qiu R, Kitzman JO, Shendure J. Chromosome-scale scaffolding of de novo genome assemblies based on chromatin interactions. Nat Biotechnol. 2013;31:1119–25. 10.1038/nbt.2727.

67. Zimin AV, Salzberg SL. The genome polishing tool POLCA makes fast and accurate corrections in genome assemblies. PLOS Comput Biol. 2020;16:e1007981. 10.1371/journal.pcbi.1007981.

68. Law CJ, Slater GJ, Mehta RS. Lineage Diversity and Size Disparity in Musteloidea: Testing Patterns of Adaptive Radiation Using Molecular and Fossil-Based Methods. Syst Biol. 2018;67:127–44. 10.1093/sysbio/syx047.

69. Hassanin A, Veron G, Ropiquet A, Vuuren BJ van, Lécu A, Goodman SM, et al. Evolutionary history of Carnivora (Mammalia, Laurasiatheria) inferred from mitochondrial genomes. PLOS ONE. 2021;16:e0240770. 10.1371/journal.pone.0240770.

70. Rozhnov VV, Meschersky IG, Pishchulina SL, Simakin LV. Genetic analysis of sable (Martes zibellina) and pine marten (M. martes) populations in sympatric part of distribution area in the northern Urals. Russ J Genet. 2010;46:488–92. 10.1134/S1022795410040150.

71. Kassal BYu, Sidorov GN. Distribution of the sable (Martes zibellina) and the pine marten (Martes martes) in Omsk oblast and biogeographic effects of their hybridization. Russ J Biol Invasions. 2013;4:105–15. 10.1134/S2075111713020070.

