## Supplementary Figures and Tables for "Novel chromosome-length genome assemblies of three distinct subspecies of pine marten, sable, and yellow-throated marten (genus *Martes*, family Mustelidae)"

**Supplementary figure SF1.** The distribution of 23-mers and the estimated genome sizes for *M. z. zibellina* (10xmzib), *M. m. uralensis* (10xmmar) and *M. fl. aterrima* (10xmfla).

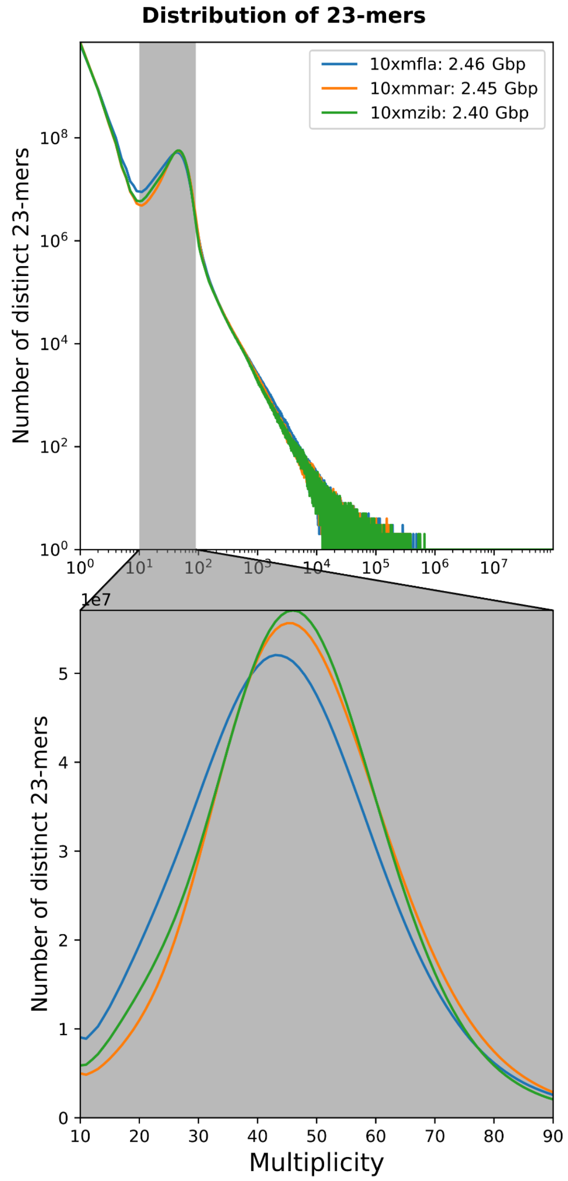

**Supplementary figure SF2.** Aligned G-banded chromosomes of five marten subspecies, only showing chr11, chr12, chr15, and chr18. Black dots indicate positions of centromeres

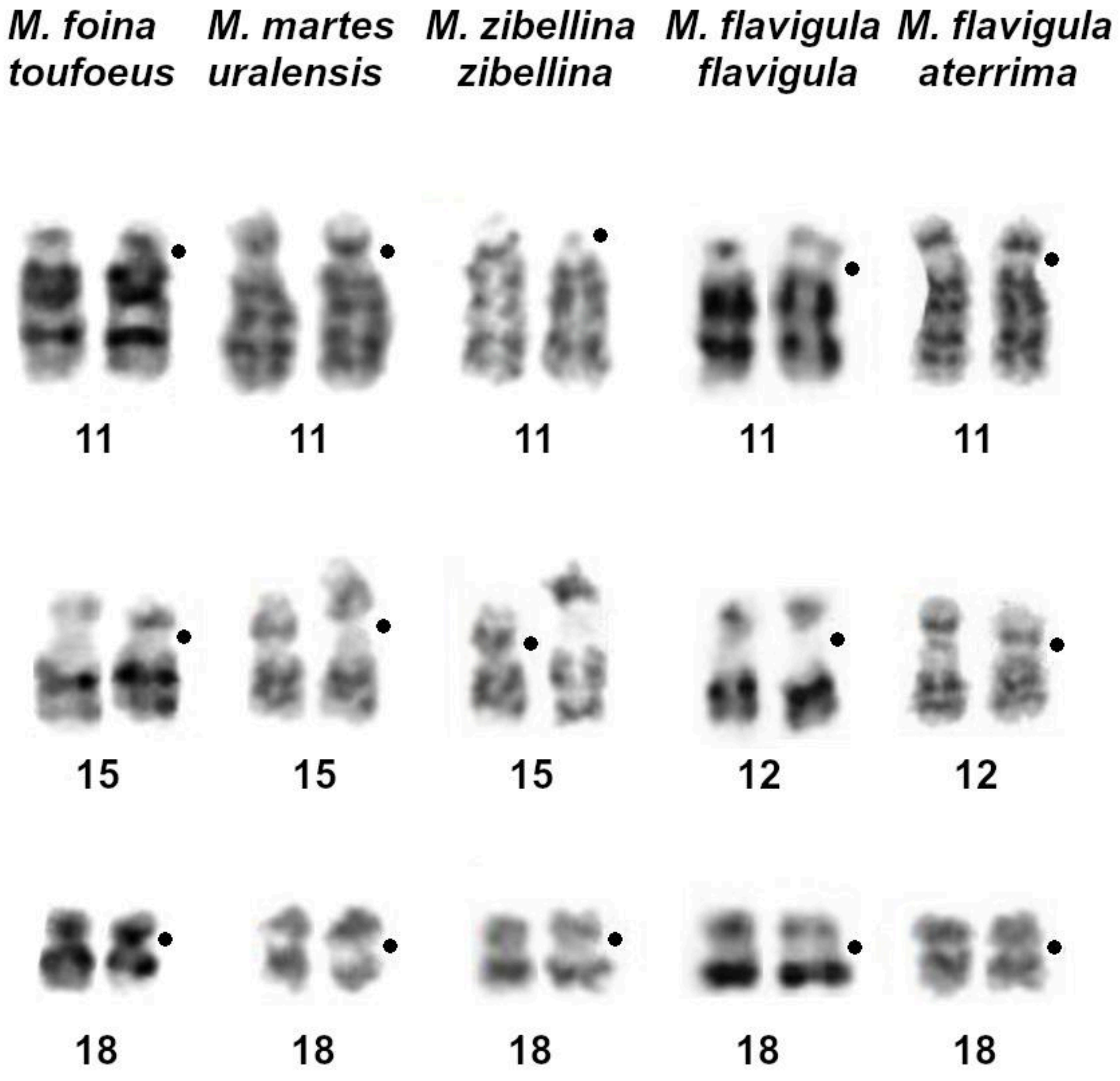

**Supplementary figure SF3.** Inverted p-arm of chr11 in the assembly of *M. fl. flavigula*

The original chromosome is shown on the left and the corrected chromosome is shown on the right. . RC - reverse complement

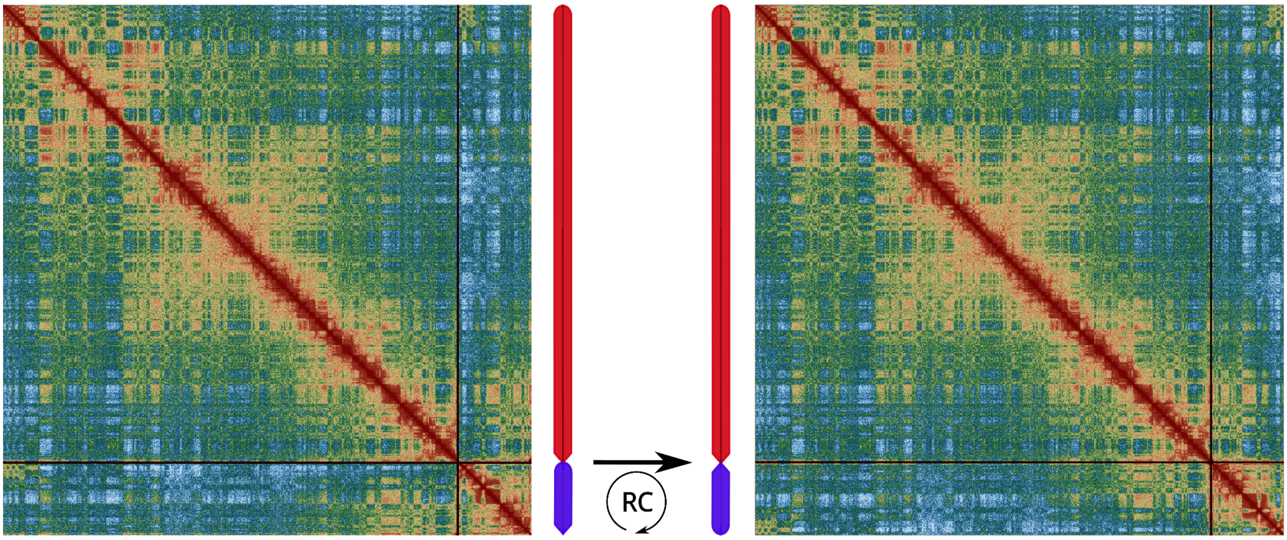


**Supplementary figure SF4.** Telomeric merge of the chromosomal arms of chr12 in the assembly of *M. fl. flavigula*

The original chromosome is shown on the left and the corrected chromosome is shown on the right. . RC - reverse complement.

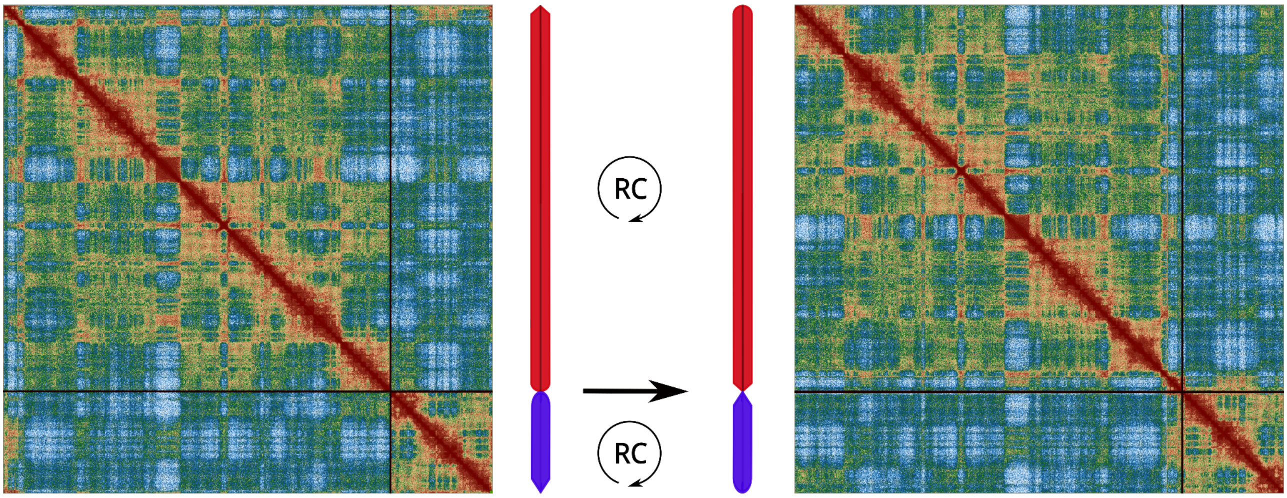

**Supplementary figure SF5.** Telomeric merge of the chromosomal arms of chr18 in the assembly of *M. fl. flavigula*

The original chromosome is shown on the left and the corrected chromosome is shown on the right. . RC - reverse complement.

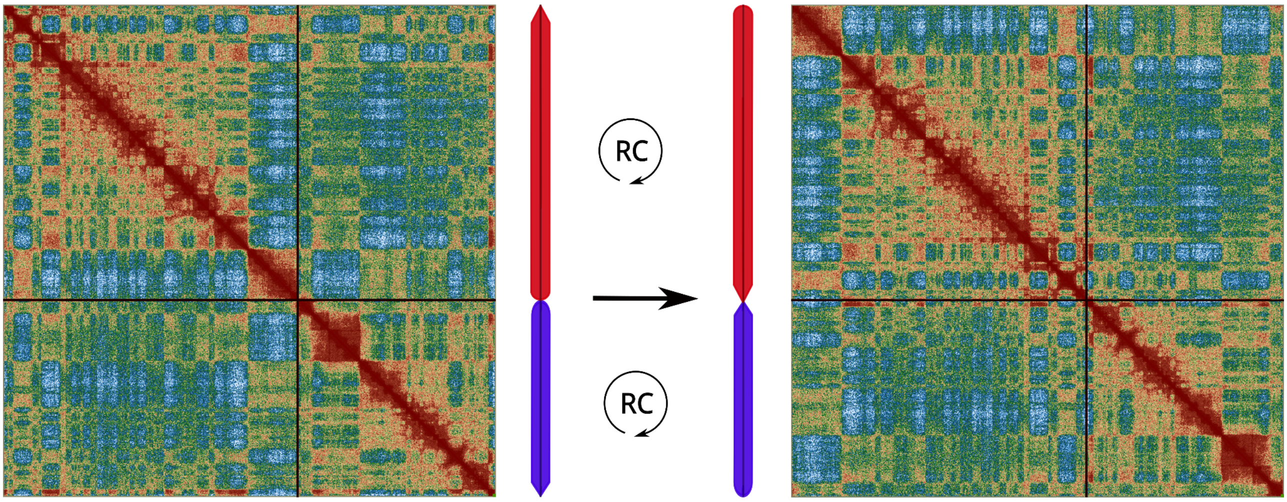

**Supplementary figure SF6.** Telomeric merge of the chromosomal arms of chr15 in the assembly of *M. fl. flavigula* (fine scale).

A, B, C - three contigs of chr 18. ABC - order of the contigs in the assembly, B' A' C - corrected order and orientation of the contigs. RC - reverse complement.

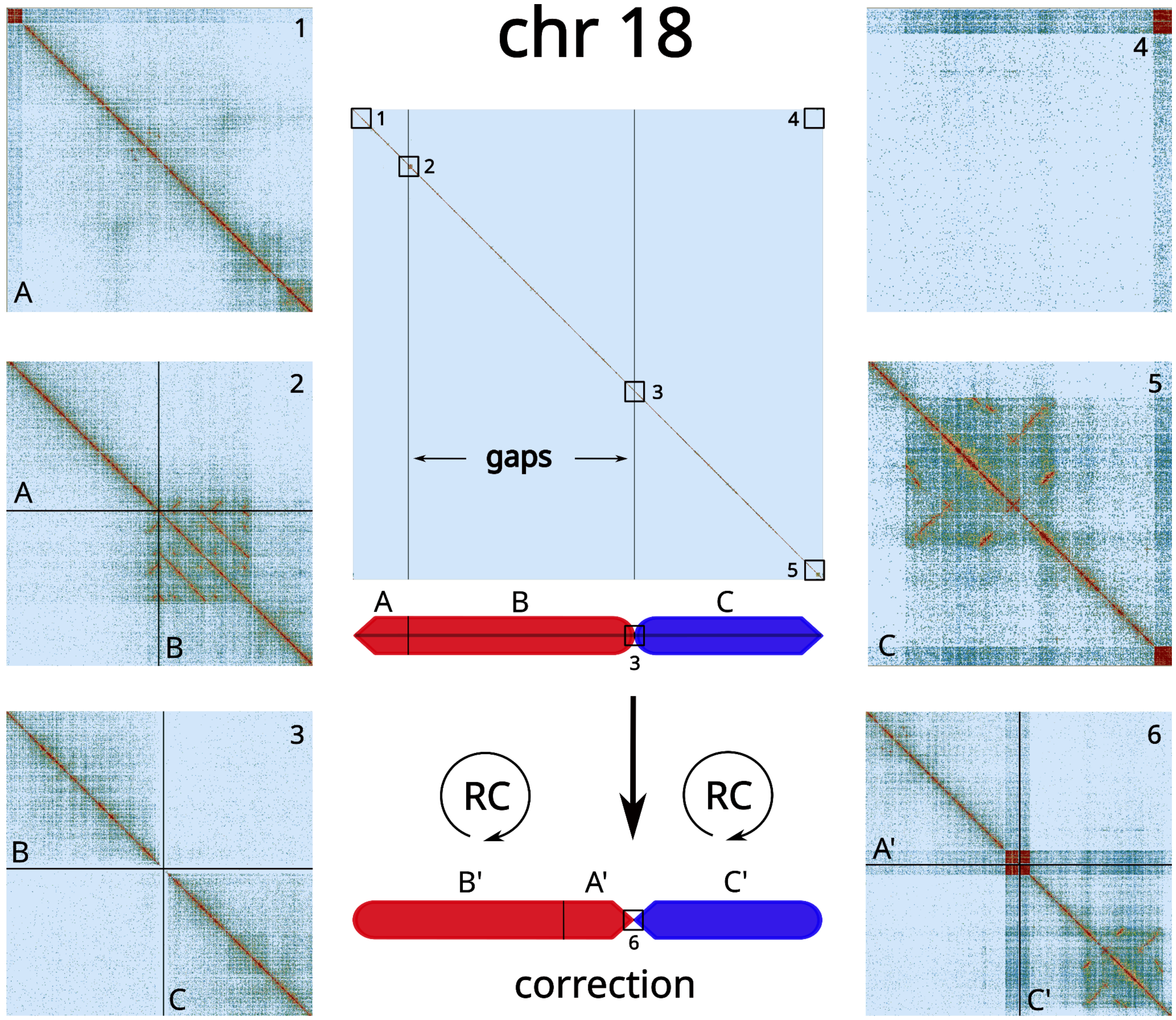


**Supplementary figure SF7.** Telomeric merge of the chromosomal arms of chr15 in the assembly of *M. f. foina*

The original chromosome is shown on the left, the corrected chromosome is shown on the right. . RC - reverse complement.

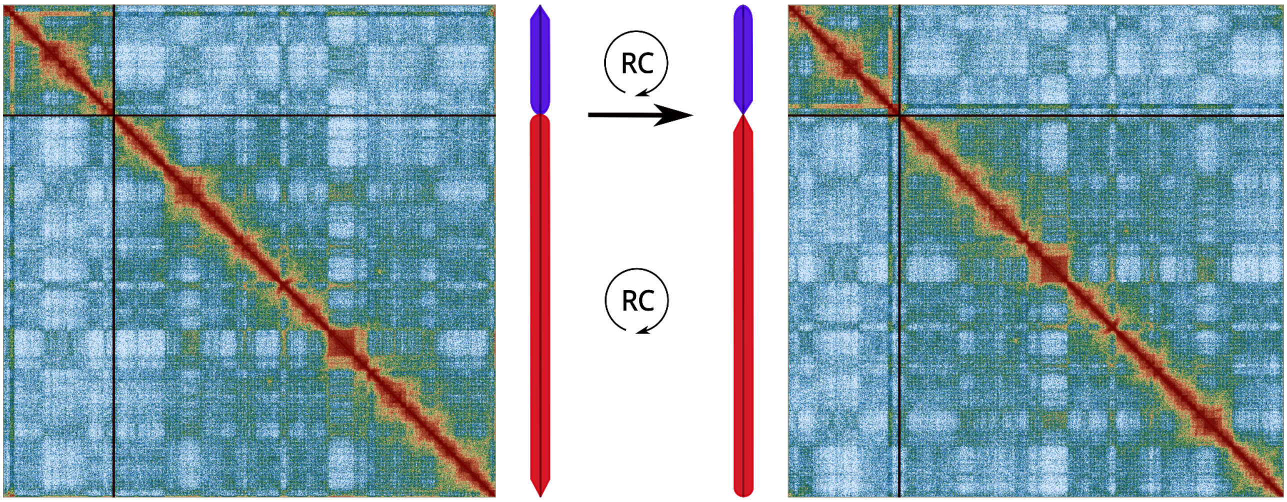


**Supplementary Figure SF8** *M. fl. aterrima* PAR coverage.

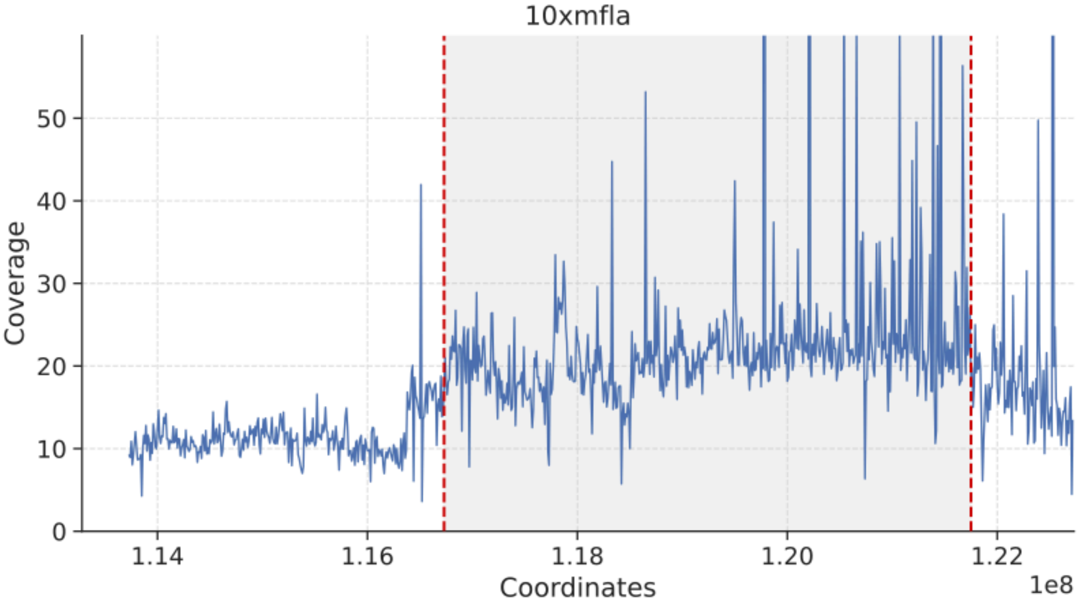


**Supplementary Figure SF9**. Overlap of the named genes in the annotations of *M. z. zibellina*, *M. m. uralensis* and *M. fl. flavigula* genome assemblies.

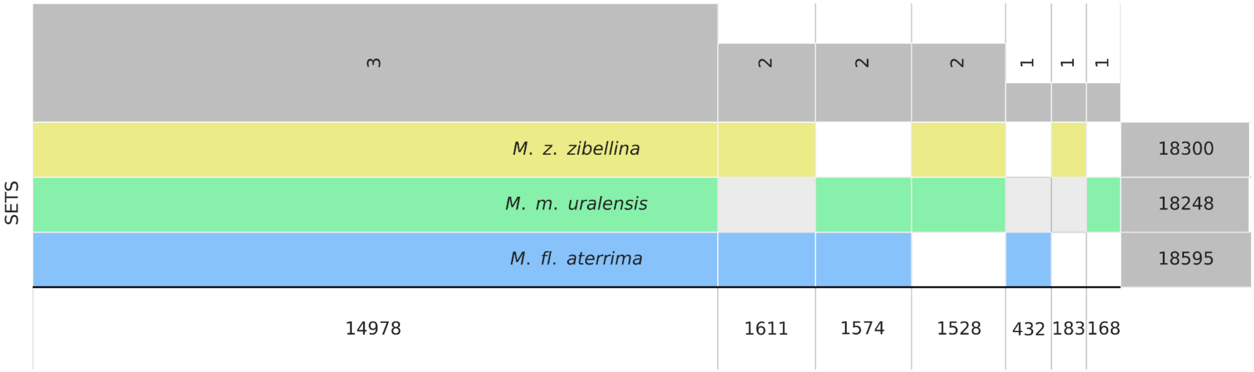

#### Supplementary Tables:

**Supplementary Table ST1**. Described subspecies of the eight recognized marten species.

| Species | Common name | Subspecies* |
|---|---|---|
| *Martes zibellina (17)* | Sable | *M. z. zibellina* Linnaeus, 1758 (syn.: *alba, asiatica, fusco-flavescens, maculata, ochracea, rupestris, sylvestris*)<br>*M. z. princeps* Birula, 1922 (syn.: *baicalensis, vitimensis*)<br>*M. z. angarensis* Timofeev et Nadeev, 1955<br>*M. z. arsenjevi* Bobrinskii, Kuznetsov et Kuzyakin, 1944<br>*M. z. averini* Bashanov, 1943 (syn.: *altaica, jurgensoni*)<br>*M. z. brachyura* Temmink, 1844<br>*M. z. ilimpiensis* Timofeev et Nadeev, 1955<br>*M. z. jakutensis* Novikov, 1956<br>*M. z. kamtschadalica* Birula, 1918 (syn.: *kamtschatica*)<br>*M. z. linkouensis* Ma et Wu, 1981<br>*M. z. obscura* Timofeev et Nadeev, 1955<br>*M. z. sahalinensis* Ognev, 1925<br>*M. z. sajanensis* Ognev, 1925<br>*M. z. schantaricus* Bobrinskii, Kuznetsov et Kuzyakin, 1944<br>*M. z. tomensis* Timofeev et Nadeev, 1955<br>*M. z. tungussensis* Kuznetsov, 1941<br>*M. z. yeniseensis* Ognev, 1925 |
| *Martes martes (10)* | Pine marten | *M. m. martes* Linnaeus, 1758<br>*M. m. uralensis* Kuznetzov, 1941<br>*M. m. latinorum* Barrett-Hamilton, 1904<br>*M. m. notialis* Cavazza, 1912<br>*M. m. lorenzi* Ognev, 1926<br>*M. m. ruthena* Ognev, 1926<br>*M. m. borealis* Kuznetzov, 1941<br>*M. m. sabaneevi* Jurgenson, 1947<br>*M. m. kuznetsovi* Pavlinov et Rossolimo, 1987<br>*M. m. minoricensis* Alcover, Delibes, Gosálbez et Nadal, 1986 |
| *Martes foina (11)* | Stone marten | *M. f. foina* Erxleben, 1777<br>*M. f. bosniaca* Brass, 1911<br>*M. f. bunites* Bate, 190<br>*M. f. intermedia* Severtzov, 1873<br>*M. f. kozlovi* Ognev, 1931<br>*M. f. mediterranea* Barrett-Hamilton, 1898<br>*M. f. milleri* Festa, 1914<br>*M. f. nehringi* Satunin, 1906<br>*M. f. rosanowi* Martino and Martino, 1917<br>*M. f. syriaca* Nehring, 1902<br>*M. f. toufoeus* Hodgson, 1842 |
| *Martes melampus (3)* | Japanese marten | *M. m. melampus* Wagner, 1840<br>*M. m. tsuensis* Thomas, 1897<br>*M. m. coreensis* Kuroda et Mori, 1923 |

| | | |
|---|---|---|
| *Martes americana (6)* | American marten | *M. a. americana* Turton, 1806<br>*M. a. abieticola* Preble, 1902<br>*M. a. abietinoides* Gray, 1865<br>*M. a. actuosa* Osgood, 1900<br>*M. a. atrata* Bangs, 1897 (syn.: *brumalis*)<br>*M. a. kenaiensis* Elliot, 1903 |
| *Martes caurina (6)* | Pacific marten | *M. c. caurina* Merriam, 1890<br>*M. c. humboldtensis* Grinnell et Dixon, 1926<br>*M. c. origenes* Rhoads, 1902<br>*M. c. sierrae* Grinnell and Storer, 1916<br>*M. c. vancouverensis* Grinnell et Dixon, 1926<br>*M. c. nobilis* Hall, 1926 |
| *Martes (Charronia) flavigula (10)* | Yellow-throated marten | *M. (Ch.) fl. aterrima* Pallas, 1811 (syn.: *borealis*)<br>*M. (Ch.) fl. flavigula* Boddaert, 1785<br>*M. (Ch.) fl. chrisospila* Swinhoe, 1866<br>*M. (Ch.) fl. peninsularis* Bonhote, 1901<br>*M. (Ch.) fl. indochinensis* Kloss, 1916<br>*M. (Ch.) fl. saba* Chasen et Kloss, 1931<br>*M. (Ch.) fl. chrysospila* Swinhoe, 1866 (syn.: *xanthospila*)<br>*M. (Ch.) fl. hainana* Hsu et Wu, 1981<br>*M. (Ch.) fl. henrici* Schinz, 1845<br>*M. (Ch.) fl. robinsoni* Pocock, 1936 |
| *Martes (Charronia) gwatkinsii* | Nilgiri yellow-throated marten | Monotypic |

* number of subspecies varies depending on the source

**Supplementary Table ST2**. Source and information of the RNA-seq data used for gene annotation of the new marten genome assemblies reported in this study.

| Species | ID | Instrument | Source | Tissue | Source |
|---|---|---|---|---|---|
| *M. martes* | ERR11872609 | Illumina NovaSeq 6000 | United Kingdom | liver | (O'Brien and Januszczak 2024) |
| *M. zibellina* | SRR13013010 | Illumina HiSeq 2500 | China | adipose | (Xia et al. 2021) |
| *M. zibellina* | SRR13013011 | Illumina HiSeq 2500 | China | adipose | (Xia et al. 2021) |
| *M. zibellina* | SRR8074161 | Illumina HiSeq 2500 | China, Greater Khingan Mountains | spleen | (Liu et al. 2020) |
| *M. zibellina* | SRR8074163 | Illumina HiSeq 2500 | China, Greater Khingan Mountains | lung | (Liu et al. 2020) |
| *M. zibellina* | SRR8074165 | Illumina HiSeq 2500 | China, Greater Khingan Mountains | muscle | (Liu et al. 2020) |
| *M. zibellina* | SRR8074168 | Illumina HiSeq 2500 | China, Greater Khingan Mountains | kidney | (Liu et al. 2020) |
| *M. zibellina* | SRR8074169 | Illumina HiSeq 2500 | China, Greater Khingan Mountains | heart | (Liu et al. 2020) |

| Species | ID | Instrument | Source | Tissue | Source |
|---|---|---|---|---|---|
| *M. zibellina* | SRR31089886 | Illumina HiSeq X | Altai Krai, Complex "Magistralny" | liver | this study |
| *M. zibellina* | SRR31089885 | Illumina HiSeq X | Altai Krai, Complex "Magistralny" | hemispheres | this study |
| *M. flavigula* | SRR21460068 | Illumina NovaSeq 6000 | China, pr. Sichuan | intestine | (Mei et al. 2023) |
| *M. flavigula* | SRR21460069 | Illumina NovaSeq 6000 | China, pr. Sichuan | spleen | (Mei et al. 2023) |
| *M. flavigula* | SRR21460070 | Illumina NovaSeq 6000 | China, pr. Sichuan | heart | (Mei et al. 2023) |
| *M. flavigula* | SRR21460071 | Illumina NovaSeq 6000 | China, pr. Sichuan | pancreas | (Mei et al. 2023) |
| *M. flavigula* | SRR21460072 | Illumina NovaSeq 6000 | China, pr. Sichuan | kidney | (Mei et al. 2023) |
| *M. flavigula* | SRR21460074 | Illumina NovaSeq 6000 | China, pr. Sichuan | testes | (Mei et al. 2023) |

**Supplementary Table ST3.** Quality metrics of six marten genome assemblies. Only contigs of ≥1000 bp and longer were taken into account.

| Species | ***M. martes uralensis*** | ***M. martes martes*** | ***M. zibellina zibellina*** | ***M. zibellina princeps*** | ***M. flavigula aterrima*** | ***M. flavigula flavigula*** |
|---|---|---|---|---|---|---|
| Source | **this study** | (O'Brien and Januszczak 2024) | **this study** | (Liu et al. 2020) | **this study** | (Mei et al. 2023) |
| 2n | 38 | 38 | 38 | 38 | 40 | 40 |
| Number of chromosomal scaffolds | 19 | 19 | 19 | - | 20 | 20 |
| Number of scaffolds | 6354 | 477 | 5965 | 15682 | 22884 | 128 |
| Total length, Gbp | 2.40 | 2.48 | 2.39 | 2.42 | 2.45 | 2.45 |
| GC, % | 41.25 | 42.37 | 41.25 | 39.99 | 41.26 | 41.72 |
| Ns, Mbp | 24.95 | 0.45 | 24.89 | 105.11 | 25.9 | 0.0085 |
| N50, Mbp | 144.64 | 146.29 | 143.64 | 5.2 | 137.4 | 143.11 |
| NCBI assembly ID | GCA_040938825.1 | GCA_963455335.1 | GCA_040938815.1 | GCA_012583365.1 | GCA_040938845.1 | GCA_029410595.1 |

**Supplementary Table ST4.** BUSCO statistics of the marten assemblies. BUSCO analysis was performed using BUSCO v5.4.2 and mammalia_odb v.10 (2021-02-19).

| Species | ***M. zibellina zibellina*** | ***M. zibellina linkouensis*** | ***M. martes uralensis*** | ***M. martes martes*** | ***M. flavigula aterrima*** | ***M. flavigula flavigula*** |
|---|---|---|---|---|---|---|
| Complete BUSCOs | 8870 (96.1%) | 8742 (94.8%) | 8895 (96.4%) | 8892 (96.3%) | 8636 (93.6%) | 8936 (96.9%) |
| Complete and single-copy BUSCOs | 8816 (95.6%) | 8679 (94.1%) | 8843 (95.8%) | 8833 (95.7%) | 8501 (92.1%) | 8874 (96.2%) |

| **Species** | ***M. zibellina zibellina*** | ***M. zibellina linkouensis*** | ***M. martes uralensis*** | ***M. martes martes*** | ***M. flavigula aterrima*** | ***M. flavigula flavigula*** |
|---|---|---|---|---|---|---|
| Complete and duplicated BUSCOs | 54 (0.6%) | 63 (0.7%) | 52 (0.6%) | 59 (0.6%) | 135 (1.5%) | 62 (0.7%) |
| Fragmented BUSCOs | 100 (1.1%) | 150 (1.6%) | 86 (0.9%) | 82 (0.9%) | 195 (2.1%) | 60 (0.7%) |
| Missing BUSCOs | 256 (2.8%) | 334 (3.6%) | 245 (2.7%) | 252 (2.8%) | 395 (4.3%) | 230 (2.4%) |

**Supplementary Table ST5.** Chromosome assignments.

| *M. zibellina zibellina* | | | *M. martes uralensis* | | | *M. martes martes* | | *M. foina toufoeus** | | | *M. foina foina* | | *M. flavigula aterrima* | | | *M. flavigula flavigula* | |
|---|---|---|---|---|---|---|---|---|---|---|---|---|---|---|---|---|---|
| **Scaffold ID, DNAzoo** | **Scaffold ID, NCBI** | **Chromosome** | **Scaffold ID, DNAzoo** | **Scaffold ID, NCBI** | **Chromosome** | **Scaffold ID, NCBI** | **Chromosome** | **Scaffold ID, DNAzoo** | **Scaffold ID, NCBI** | **Chromosome** | **Scaffold ID, NCBI** | **Chromosome** | **Scaffold ID, DNAzoo** | **Scaffold ID, NCBI** | **Chromosome**** | **Scaffold ID, NCBI** | **Chromosome**** |
| HiC_scaffold_2 | CM081924.1 | chr1 | HiC_scaffold_2 | CM081865.1 | chr1 | OY734064.1 | chr1 | HiC_scaffold_2 | CM081884.1 | chr1 | OZ199605.1 | chr1 | HiC_scaffold_2 | CM081904.1 | chr1 | CM055620.1 | chr1 |
| HiC_scaffold_1 | CM081925.1 | chr2 | HiC_scaffold_1 | CM081866.1 | chr2 | OY734063.1 | chr2 | HiC_scaffold_1 | CM081885.1 | chr2 | OZ199604.1 | chr2 | HiC_scaffold_1 | CM081905.1 | chr2 | CM055619.1 | chr2 |
| HiC_scaffold_3 | CM081926.1 | chr3 | HiC_scaffold_3 | CM081867.1 | chr3 | OY734065.1 | chr3 | HiC_scaffold_3 | CM081886.1 | chr3 | OZ199606.1 | chr3 | HiC_scaffold_3 | CM081906.1 | chr3 | CM055621.1 | chr3 |
| HiC_scaffold_4 | CM081927.1 | chr4 | HiC_scaffold_4 | CM081868.1 | chr4 | OY734066.1 | chr4 | HiC_scaffold_4 | CM081887.1 | chr4 | OZ199607.1 | chr4 | HiC_scaffold_4 | CM081907.1 | chr4 | CM055622.1 | chr4 |
| HiC_scaffold_5 | CM081928.1 | chr5 | HiC_scaffold_5 | CM081869.1 | chr5 | OY734067.1 | chr5 | HiC_scaffold_5 | CM081888.1 | chr5 | OZ199608.1 | chr5 | HiC_scaffold_5 | CM081908.1 | chr5 | CM055623.1 | chr5 |
| HiC_scaffold_6 | CM081929.1 | chr6 | HiC_scaffold_6 | CM081870.1 | chr6 | OY734068.1 | chr6 | HiC_scaffold_6 | CM081889.1 | chr6 | OZ199609.1 | chr6 | HiC_scaffold_6 | CM081909.1 | chr6 | CM055624.1 | chr6 |
| HiC_scaffold_7 | CM081930.1 | chr7 | HiC_scaffold_7 | CM081871.1 | chr7 | OY734069.1 | chr7 | HiC_scaffold_7 | CM081890.1 | chr7 | OZ199610.1 | chr7 | HiC_scaffold_9 | CM081912.1 | chr9 | CM055627.1 | chr9 |
| - | - | - | - | - | - | - | - | | - | - | - | - | HiC_scaffold_19 | CM081922.1 | chr19 | CM055637.1 | chr19 |
| HiC_scaffold_9 | CM081931.1 | chr8 | HiC_scaffold_9 | CM081872.1 | chr8 | OY734071.1 | chr8 | HiC_scaffold_9 | CM081891.1 | chr8 | OZ199612.1 | chr8 | HiC_scaffold_8 | CM081911.1 | chr8 | CM055626.1 | chr8 |
| HiC_scaffold_8 | CM081932.1 | chr9 | HiC_scaffold_8 | CM081873.1 | chr9 | OY734070.1 | chr9 | HiC_scaffold_8 | CM081892.1 | chr9 | OZ199611.1 | chr9 | HiC_scaffold_7 | CM081910.1 | chr7 | CM055625.1 | chr7 |
| HiC_scaffold_10 | CM081933.1 | chr10 | HiC_scaffold_10 | CM081874.1 | chr10 | OY734073.1 | chr10 | HiC_scaffold_10 | CM081893.1 | chr10 | OZ199614.1 | chr10 | HiC_scaffold_10 | CM081913.1 | chr10 | CM055628.1 | chr10 |
| HiC_scaffold_11 | CM081934.1 | chr11 | HiC_scaffold_11 | CM081875.1 | chr11 | OY734074.1 | chr11 | HiC_scaffold_11 | CM081894.1 | chr11 | OZ199615.1 | chr11 | HiC_scaffold_11 | CM081914.1 | chr11 | CM055629.1 | chr11 |
| HiC_scaffold_14 | CM081935.1 | chr12 | HiC_scaffold_14 | CM081876.1 | chr12 | OY734077.1 | chr12 | HiC_scaffold_14 | CM081895.1 | chr12 | OZ199618.1 | chr12 | HiC_scaffold_14 | CM081916.1 | chr13 | CM055632.1 | chr13 |
| HiC_scaffold_12 | CM081936.1 | chr13 | HiC_scaffold_12 | CM081877.1 | chr13 | OY734076.1 | chr13 | HiC_scaffold_12 | CM081896.1 | chr13 | OZ199616.1 | chr13 | HiC_scaffold_12 | CM081918.1 | chr15 | CM055630.1 | chr15 |
| HiC_scaffold_13 | CM081937.1 | chr14 | HiC_scaffold_13 | CM081878.1 | chr14 | OY734075.1 | chr14 | HiC_scaffold_13 | CM081897.1 | chr14 | OZ199617.1 | chr14 | HiC_scaffold_13 | CM081917.1 | chr14 | CM055631.1 | chr14 |
| HiC_scaffold_16 | CM081938.1 | chr15 | HiC_scaffold_16 | CM081879.1 | chr15 | OY734079.1 | chr15 | HiC_scaffold_16 | CM081898.1 | chr15 | OZ199620.1 | chr15 | HiC_scaffold_16 | CM081915.1 | chr12 | CM055634.1 | chr12 |
| HiC_scaffold_15 | CM081939.1 | chr16 | HiC_scaffold_15 | CM081880.1 | chr16 | OY734078.1 | chr16 | HiC_scaffold_15 | CM081899.1 | chr16 | OZ199619.1 | chr16 | HiC_scaffold_15 | CM081919.1 | chr16 | CM055633.1 | chr16 |
| HiC_scaffold_17 | CM081940.1 | chr17 | HiC_scaffold_17 | CM081881.1 | chr17 | OY734080.1 | chr17 | HiC_scaffold_17 | CM081900.1 | chr17 | OZ199621.1 | chr17 | HiC_scaffold_17 | CM081920.1 | chr17 | CM055635.1 | chr17 |
| HiC_scaffold_18 | CM081941.1 | chr18 | HiC_scaffold_18 | CM081882.1 | chr18 | OY734081.1 | chr18 | HiC_scaffold_18 | CM081901.1 | chr18 | OZ199622.1 | chr18 | HiC_scaffold_18 | CM081921.1 | chr18 | CM055636.1 | chr18 |
| HiC_scaffold_19 | CM081942.1 | chrX | HiC_scaffold_19 | CM081883.1 | chrX | OY734072.1 | chrX | HiC_scaffold_19 | CM081902.1 | chrX | OZ199613.1 | chrX | HiC_scaffold_20 | CM081923.1 | chrX | CM055638.1 | chrX |

** chromosome nomenclature from (Tomarovsky et al. 2025)*

*** - chromosome nomenclature of the M. flavigula aterrima is different from other martens*

**Supplementary Table ST6.** Interspersed repeat content in the new and previously published marten genome assemblies.

| Transposable elements | *M. zibellina zibellina* | | *M. zibellina princeps* | | *M. martes uralensis* | | *M. martes martes* | | *M. flavigula aterrima* | | *M. flavigula flavigula* | | *M. foina toufoeus* | | *M. foina foina* | |
|---|---|---|---|---|---|---|---|---|---|---|---|---|---|---|---|---|
| | **Length, bp** | **Length, %** | **Length, bp** | **Length, %** | **Length, bp** | **Length, %** | **Length, bp** | **Length, %** | **Length, bp** | **Length, %** | **Length, bp** | **Length, %** | **Length, bp** | **Length, %** | **Length, bp** | **Length, %** |
| **Retroelements** | 870’871’585 | 36.37 | 824’279’833 | 34.05 | 878’539’291 | 36.54 | 899’003’202 | 36.18 | 912’279’015 | 36.86 | 918’790’523 | 37.52 | 875’539’264 | 36.19 | 876’077’360 | 36.68 |
| SINEs | 237’444’556 | 9.92 | 228’318’320 | 9.43 | 238’992’690 | 9.94 | 241’509’745 | 9.72 | 246’594’632 | 9.96 | 245’280’198 | 10.02 | 238’158’404 | 9.84 | 238’216’477 | 9.97 |
| Penelope | 57’707 | 0 | 57’078 | 0 | 56’129 | 0 | 56’341 | 0 | 55’804 | 0 | 57’379 | 0 | 56’066 | 0 | 55’598 | 0 |
| LINEs | 518’969’807 | 21.67 | 482’489’753 | 19.93 | 524’763’476 | 21.83 | 540’808’650 | 21.77 | 546’947’504 | 22.1 | 555’589’110 | 22.69 | 521’971’290 | 21.57 | 522’867’180 | 21.89 |
| CRE/SLACS | 0 | 0 | 0 | 0 | 0 | 0 | 0 | 0 | 0 | 0 | 0 | 0 | 0 | 0 | 0 | 0 |
| L2/CR1/Rex | 88’603’222 | 3.7 | 88’307’113 | 3.65 | 88’635’894 | 3.69 | 89’047’902 | 3.58 | 88’662’032 | 3.58 | 88’710’630 | 3.62 | 88’385’088 | 3.65 | 88’622’296 | 3.71 |
| R1/LOA/Jockey | 0 | 0 | 0 | 0 | 0 | 0 | 0 | 0 | 0 | 0 | 0 | 0 | 0 | 0 | 0 | 0 |
| R2/R4/NeSL | 83’901 | 0 | 85’105 | 0 | 82’868 | 0 | 82’455 | 0 | 82’828 | 0 | 81’710 | 0 | 85’171 | 0 | 83’847 | 0 |
| RTE/Bov-B | 2’875’393 | 0.12 | 2’870’448 | 0.12 | 2’879’106 | 0.12 | 2’871’782 | 0.12 | 2’876’245 | 0.12 | 2’860’979 | 0.12 | 2’859’653 | 0.12 | 2’854’379 | 0.12 |
| L1/CIN4 | 427’361’651 | 17.85 | 391’181’748 | 16.16 | 433’119’245 | 18.01 | 448’760’664 | 18.06 | 455’280’984 | 18.39 | 463’890’630 | 18.94 | 430’594’586 | 17.8 | 431’260’601 | 18.05 |
| **LTR elements** | 114’457’222 | 4.78 | 113’471’760 | 4.69 | 114’783’125 | 4.77 | 116’684’807 | 4.7 | 118’736’879 | 4.8 | 117’921’215 | 4.82 | 115’409’570 | 4.77 | 114’993’703 | 4.81 |
| BEL/Pao | 0 | 0 | 0 | 0 | 0 | 0 | 0 | 0 | 0 | 0 | 0 | 0 | 0 | 0 | 0 | 0 |
| Ty1/Copia | 0 | 0 | 0 | 0 | 0 | 0 | 0 | 0 | 0 | 0 | 0 | 0 | 0 | 0 | 0 | 0 |
| Gypsy/DIRS1 | 3’450’189 | 0.14 | 3’449’216 | 0.14 | 3’459’917 | 0.14 | 3’508’652 | 0.14 | 3’443’291 | 0.14 | 3’453’002 | 0.14 | 3’425’117 | 0.14 | 3’442’695 | 0.14 |
| Retroviral | 108’815’544 | 4.54 | 107’841’622 | 4.46 | 109’130’344 | 4.54 | 110’974’522 | 4.47 | 113’099’281 | 4.57 | 112’282’421 | 4.59 | 109’791’419 | 4.54 | 109’372’470 | 4.58 |
| **DNA transposons** | 69076’893 | 2.88 | 68’716’294 | 2.84 | 69’092’488 | 2.87 | 69’427’432 | 2.79 | 69’741’546 | 2.82 | 69’327’291 | 2.83 | 69’012’541 | 2.85 | 69’030’833 | 2.89 |
| hobo-Activator | 48’344’438 | 2.02 | 48’069’960 | 1.99 | 48’360’389 | 2.01 | 48’600’897 | 1.96 | 48’800’534 | 1.97 | 48’514’362 | 1.98 | 48’291’179 | 2 | 48’325’200 | 2.02 |
| Tc1-IS630-Pogo | 19’325’274 | 0.81 | 19’239’567 | 0.79 | 19’326’908 | 0.8 | 19’418’533 | 0.78 | 19’532’221 | 0.79 | 19’411’325 | 0.79 | 19’322’784 | 0.8 | 19’304’756 | 0.81 |
| En-Spm | 0 | 0 | 0 | 0 | 0 | 0 | 0 | 0 | 0 | 0 | 0 | 0 | 0 | 0 | 0 | 0 |
| MULE-MuDR | 65’089 | 0 | 65’028 | 0 | 61’858 | 0 | 64’504 | 0 | 62’517 | 0 | 62’108 | 0 | 58’406 | 0 | 62’634 | 0 |
| PiggyBac | 198’772 | 0.01 | 198’170 | 0.01 | 200’754 | 0.01 | 201’573 | 0.01 | 203’349 | 0.01 | 200’518 | 0.01 | 199’207 | 0.01 | 200’766 | 0.01 |
| Tourist/Harbinger | 62’378 | 0 | 62’916 | 0 | 63’052 | 0 | 62’292 | 0 | 61’914 | 0 | 61’988 | 0 | 61’753 | 0 | 62’105 | 0 |

| Transposable elements | *M. zibellina zibellina* | | *M. zibellina princeps* | | *M. martes uralensis* | | *M. martes martes* | | *M. flavigula aterrima* | | *M. flavigula flavigula* | | *M. foina toufoeus* | | *M. foina foina* | |
|---|---|---|---|---|---|---|---|---|---|---|---|---|---|---|---|---|
| | **Length, bp** | **Length, %** | **Length, bp** | **Length, %** | **Length, bp** | **Length, %** | **Length, bp** | **Length, %** | **Length, bp** | **Length, %** | **Length, bp** | **Length, %** | **Length, bp** | **Length, %** | **Length, bp** | **Length, %** |
| Other | 0 | 0 | 0 | 0 | 0 | 0 | 0 | 0 | 0 | 0 | 0 | 0 | 0 | 0 | 0 | 0 |
| **Rolling-circles** | 342’852 | 0.01 | 340’092 | 0.01 | 340’629 | 0.01 | 344’191 | 0.01 | 357’197 | 0.01 | 350’741 | 0.01 | 334’299 | 0.01 | 336’081 | 0.01 |
| **Unclassified** | 544’264 | 0.02 | 541’446 | 0.02 | 544’192 | 0.02 | 542’347 | 0.02 | 551’664 | 0.02 | 546’624 | 0.02 | 538’075 | 0.02 | 538’153 | 0.02 |
| **Total interspersed repeats** | 940’550’449 | 39.28 | 893’594’651 | 36.92 | 948’232’100 | 39.44 | 969’029’322 | 39 | 982’628’029 | 39.7 | 988’721’817 | 40.38 | 945’145’946 | 39.07 | 945’701’944 | 39.59 |
| **Small RNA** | 175’718’980 | 7.34 | 166’619’450 | 6.88 | 177’343’940 | 7.38 | 179’577’373 | 7.23 | 184’536’005 | 7.46 | 183’323’808 | 7.49 | 176’650’893 | 7.3 | 176’489’984 | 7.39 |
| **Satellites** | 0 | 0 | 0 | 0 | 0 | 0 | 0 | 0 | 0 | 0 | 0 | 0 | 0 | 0 | 0 | 0 |
| **Simple repeats** | 30’352’239 | 1.27 | 27’641’369 | 1.14 | 30’480’662 | 1.27 | 31’885’789 | 1.28 | 30’857’877 | 1.25 | 30’910’005 | 1.26 | 31’282’371 | 1.29 | 32’021’844 | 1.34 |
| **Low complexity** | 5’544’570 | 0.23 | 5’384’524 | 0.22 | 5’614’311 | 0.23 | 5’883’696 | 0.24 | 5’517’380 | 0.22 | 5’591’243 | 0.23 | 5’632’322 | 0.23 | 5’691’032 | 0.24 |

**Supplementary Table ST7**. Metrics of the gene prediction and functional annotation for the three marten genome assemblies reported in this study.

| Species | Number of gene models | Number of transcripts | Number of named genes | Complete BUSCOs (%) |
|---|---|---|---|---|
| *M. z. zibellina* | 20158 | 30837 | 18300 | 97,7% |
| *M. m. uralensis* | 20259 | 31116 | 18248 | 97,7% |
| *M. fl. aterrima* | 20521 | 30087 | 18595 | 95,1% |

### Literature:


Liu G, Zhao C, Xu D, Zhang Huanxin, Monakhov V, Shang S, Gao X, Sha W, Ma J, Zhang W, et al. 2020. First Draft Genome of the Sable, Martes zibellina. *Genome Biol. Evol.* 12:59–65.

Mei X, Liu G, Yan J, Zhao C, Wang X, Zhou S, Wei Q, Zhao S, Liu Z, Sha W, et al. 2023. A chromosome-level genome assembly of the yellow-throated marten (Martes flavigula). *Sci. Data* 10:216.

O'Brien D, Januszczak I. 2024. The genome sequence of the European pine marten, Martes martes (Linnaeus, 1758). *Wellcome Open Res.* 9.

Tomarovsky A, Khan R, Dudchenko O, Totikov A, Serdyukova NA, Weisz D, Vorobieva NV, Bulyonkova T, Abramov AV, Nie W, et al. 2025. Chromosome-length genome assembly of the stone marten (Martes foina, Mustelidae): A new view on one of the cornerstones in carnivore cytogenetics. *J. Hered.* 116:548–557.

Xia T, Zhang L, Sun G, Yang X, Zhao C, Zhang H. 2021. Insights into cold tolerance in sable (Martes zibellina) from the adaptive evolution of lipid metabolism. *Mamm. Biol.* 101:861–870.
